# Structure and dynamics of the quaternary *hunchback* mRNA translation repression complex

**DOI:** 10.1101/2020.09.08.287060

**Authors:** Jakub Macošek, Bernd Simon, Johanna-Barbara Linse, Sophie Winter, Jaelle Foot, Kathryn Perez, Mandy Rettel, Miloš T. Ivanović, Pawel Masiewicz, Brice Murciano, Mikhail M. Savitski, Jochen S. Hub, Frank Gabel, Janosch Hennig

**Affiliations:** Structural and Computational Biology Unit, European Molecular Biology Laboratory Heidelberg, Heidelberg, 69117, Germany; Biozentrum, University of Basel, Basel, 4056, Switzerland; Theoretical Physics and Center for Biophysics, Saarland University, Saarbrücken, 66123, Germany; Proteomics Core Facility, European Molecular Biology Laboratory Heidelberg, Heidelberg, 69117, Germany; Institut Biologie Structurale, University Grenoble Alpes, CEA, CNRS, Grenoble, 38044, France; Protein Expression and Purification Core Facility, European Molecular Biology Laboratory Heidelberg, Heidelberg, 69117, Germany

## Abstract

A key regulatory process during *Drosophila* development is the localized suppression of the *hunchback* mRNA translation at the posterior, which gives rise to a *hunchback* gradient governing the formation of the anterior-posterior body axis. The suppression of the RNA is achieved by a concerted action of Brain Tumour (Brat), Pumilio (Pum) and Nanos. Each protein is necessary for proper *Drosophila* development. The RNA contacts have been elucidated for the proteins individually in several atomic-resolution structures. However, the interplay of all three proteins in the RNA suppression remains a long-standing open question. We characterize the quaternary complex of the RNA-binding domains of Brat, Pum and Nanos with *hunchback* mRNA by combining NMR spectroscopy, SANS/SAXS, XL/MS with MD simulations and ITC assays. The quaternary *hunchback* mRNA suppression complex is flexible with the unoccupied nucleotides of the RNA functioning as a flexible linker between the Brat and Pum-Nanos moieties of the complex. Moreover, Brat and Pum with Nanos bind the RNA completely independently. In accordance with previous studies, showing that Brat can suppress *hunchback* mRNA independently and is distributed uniformly throughout the embryo, this suggests that *hunchback* mRNA suppression by Brat is functionally separate from the suppression by Pumilio and Nanos.

## INTRODUCTION

One of the key processes during *Drosophila* development is the formation of body axes in the embryo. This is achieved by RNA localisation and spatially restricted translation (1), which results in protein gradients along which the axes are established (2). Proper anterior-posterior axis formation is governed by localisation of two maternally transcribed genes. The first, *bicoid* mRNA, localizes to the anterior pole of the oocyte (3), whereas the second, *nanos* mRNA, localizes to the posterior pole of the oocyte (4). Translation of *bicoid* and *nanos* mRNA then results in opposing gradients of Bicoid and Nanos proteins, which in turn control translation of another maternally supplied mRNA – *hunchback* mRNA (5-8). *Hunchback* mRNA is distributed uniformly and Nanos suppresses the mRNA translation at the posterior, whereas Bicoid activates it at the anterior. The resulting anterior-posterior gradient of Hunchback then ensures proper development of abdominal and thorax structures (9,10).

Nanos is thus the *trans*-acting molecule in the posterior suppression of *hunchback* mRNA translation. The related *cis*-acting elements are located within the 3’ untranslated region (UTR) of the mRNA and are called Nanos Response Elements (NREs) (11). There are two NREs (NRE1 and NRE2), that each contain two conserved sequences called Box A (upstream) and Box B (downstream) (Figure 1A). Nanos contains a zinc finger domain (Nanos-ZnF, Figure 1B), that binds the NREs (12), but requires the *trans*acting molecules Pumilio and Brain Tumor (Brat) for suppression of *hunchback* mRNA translation (13,14). Pumilio and Nanos have been long established as *trans*-acting elements in the suppression of *hunchback* mRNA translation and a recent study revealed the underlying mechanism of their concerted recognition of *hunchback* mRNA (Figure 1B) (15). Pumilio features a Pumilio homology domain (Pum-HD), which is a sequence-specific single-stranded RNA binding domain (RBD), that binds to a site partially overlapping with Box B of NREs (16-19). In contrast, no specific motif in *hunchback* mRNA is recognized by Nanos-ZnF, which binds the RNA nevertheless with high affinity, but presumably with low sequence specificity (12,20). However, Nanos-ZnF and Pum-HD together with *hunchback* mRNA form a high affinity ternary protein-RNA complex (15). Ternary complex formation triggers RNA specificity in Nanos-ZnF for bases adjacent to the binding site of Pum-HD and further enhances the affinity of Pum-HD for its binding site. While localisation of Nanos at the posterior ensures the spatial restriction for hunchback mRNA suppression, Pumilio conveys specificity for *hunchback* mRNA binding. Thus, the concerted recognition of the RNA by Pumilio and Nanos simultaneously allows the spatially controlled suppression of *hunchback* mRNA during *Drosophila* development.

**Figure 1.**
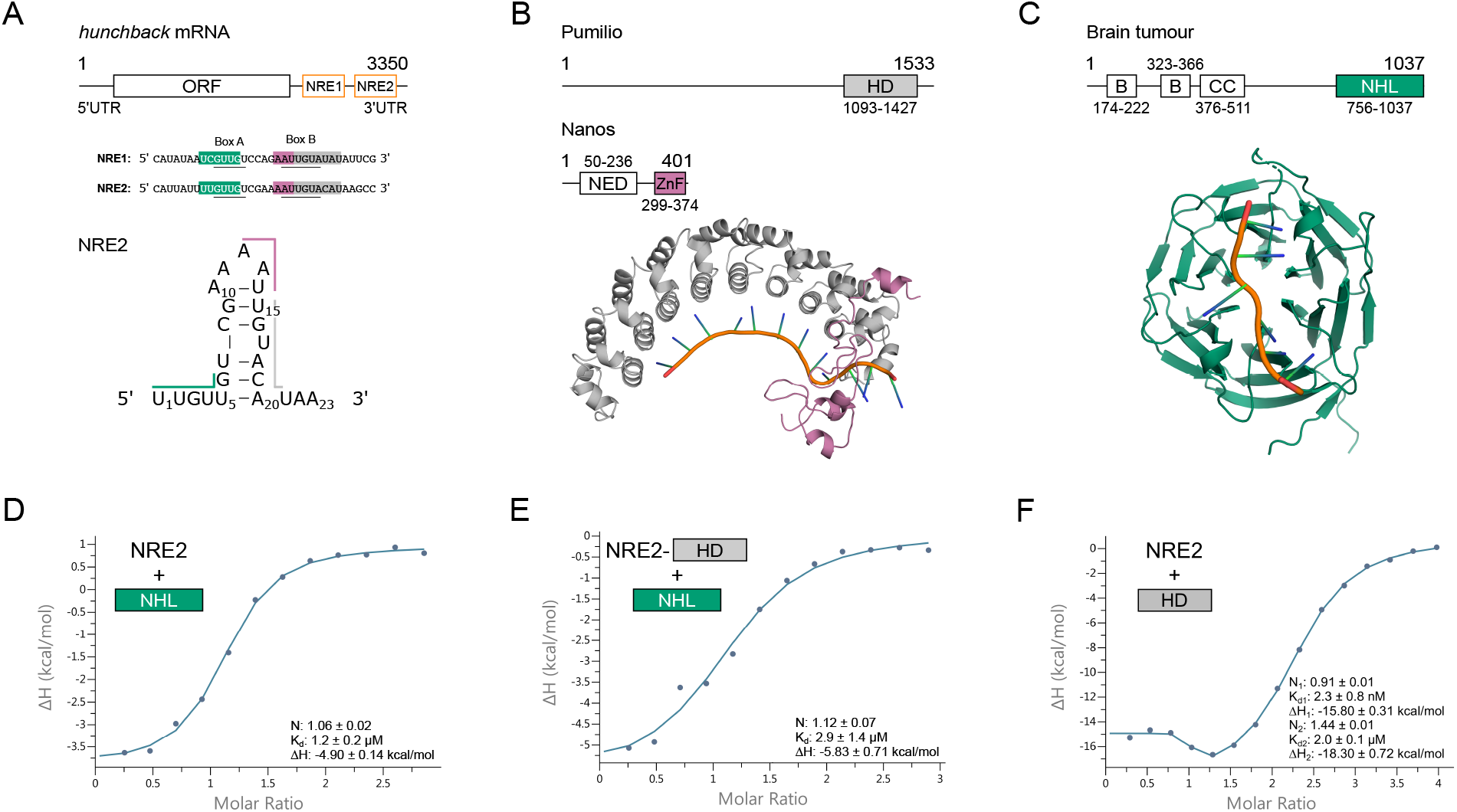
The molecules involved in suppression of *hunchback* mRNA translation and their interactions. **(A)** Two *cis*-acting elements controlling the suppression are located within the 3’ UTR and called Nanos Response Element 1 and 2 (NRE1 and NRE2). NRE2 was shown to form a stem loop (22). The *trans*acting proteins used in this study are: **(B)** the HD domain of Pumilio (Pum-HD, grey), the ZnF domain of Nanos (Nanos-ZnF, pink) and **(C)** the NHL domain of Brain tumour (Brat-NHL, green). The shading of the RNA sequences indicates the binding site of the domain matching the colour based on the previously published structures of Brat-NHL-RNA complex (PDB ID: 4zlr, middle panel) (22) and Pum HD-Nanos-ZnF-RNA complex (PDB ID: 5kl1, right panel) (15). RNA comprising NRE2 indicated in the lower part of the left panel (NRE2 RNA), the protein domains and the published high-resolution structures described above were used in this study for experimental measurements or computational modelling. **(D)** Brat-NHL-NRE2 RNA interaction studied by ITC. The measurement reveals a 1:1 interaction with a K_d_ of 1.2 ± 0.2 μM. **(E)** The interaction of Pum-HD-bound NRE2 RNA with Brat-NHL studied by ITC. A 1:1 mixture of Pum-HD and NRE2 RNA was titrated by Brat-NHL revealing a K_d_ of 2.9 ± 1.4 μM. **(F)** Pum-HD-NRE2 RNA interaction studied by ITC. The binding isotherm shows one high affinity binding event with a K_d_ of 2.3 ± 0.8 nM and one low affinity binding event with a K_d_ of 2.0 ± 0.1 μM. The first binding event consists of only 3-4 data points, but can be resolved using a shorter RNA stretch that contains only Pum-HD binding site (Supplementary Figure S2E).

Brain Tumour has been identified as another *trans*-acting molecule in the suppression of *hunchback* mRNA translation more recently (14). The NCL-1, HT2A, and LIN-41 domain of Brat (Brat-NHL) was shown to interact specifically with a motif in Box A of NREs (Figure 1C) and Brat mutants induce similar developmental defects in flies as Pumilio or Nanos mutants (21-23). Like Pumilio, Brat is uniformly distributed throughout the embryo (14), but how it contributes to the suppression of *hunchback* mRNA by Pumilio and Nanos remains unclear (24), despite a wealth of available experimental data on the relationship of Brat-NHL, Pum-HD and Nanos-ZnF. First, published data suggest that Brat, Pumilio and Nanos form a quaternary complex with *hunchback* mRNA in order to suppress its translation: in yeast four-hybrid and pull-down assays Brat-NHL, Pum-HD and Nanos-ZnF were all necessary to form a stable protein-RNA complex (14). In line with that, Pum-HD was reported to increase the affinity of Brat-NHL for *hunchback* mRNA (22). This would explain why Brat mutations induce phenotypes similar to those of Pumilio and Nanos mutants. Moreover, a vast majority of these mutants implied indirectly quaternary complex formation, as they abrogated complex formation in yeast four-hybrid assays. However, it became clear later that the point mutations substituted residues necessary for RNA binding of Brat-NHL (22), so the observed effect is likely due to abrogated RNA binding and not necessarily due to abrogation of quaternary complex formation. Furthermore, gene reporter assays in Dmel2 cells show that removing Brat-NHL has little effect on suppression of a reporter gene in the presence of Pumilio and Nanos, and that, conversely, Brat can suppress the mRNA translation independently in their absence (22,23). In summary, it remains unclear whether Brat, Pumilio and Nanos suppress hunchback mRNA translation cooperatively in a single quaternary protein-RNA complex or whether Brat suppresses *hunchback* mRNA independently of Pumilio and Nanos. To determine if the suppression activities are cooperative or independent, a combination of both functional and biophysical insights into the process is required. Since there is already a wealth of functional data, we obtained the biophysical results needed to fill the existing gap in our understanding of *hunchback* mRNA translation suppression.

We carried out an extensive structural, biophysical, and computational investigation of Pum-HD, Brat-NHL, Nanos-ZnF and their complex with *hunchback* mRNA NRE2 (*hb* complex). First, we analysed the influence of Pum-HD on the RNA binding of Brat-NHL using isothermal titration calorimetry (ITC) and showed that they do not bind RNA cooperatively. Moreover, Brat-NHL and Nanos-ZnF do not show any signs of interaction *in vitro* in the absence of RNA and no cooperativity was observed in RNA binding between Brat-NHL and Pum-HD with Nanos-ZnF. Nevertheless, Brat-NHL, Pum-HD and Nanos-ZnF assemble all on a single NRE2 in a stable stoichiometric quaternary protein-RNA complex. However, our data reveals that the complex, in context of a single NRE2 and isolated RNA binding domains, is largely flexible and that Brat-NHL in the complex moves independently of Pum-HD and Nanos-ZnF. To investigate the extent of flexibility of the complex we used unrestrained molecular dynamics (MD) simulation validated against small-angle X-ray and neutron scattering data. Collectively, our results describe a complex, in which the unoccupied nucleotides of the NRE2 RNA function as a flexible linker between the Brat-NHL and Pum-HD-Nanos-ZnF moieties of the complex, reminiscent of beads on a string. Moreover, this study highlights the importance of combining a multitude of structure analysis methods with molecular dynamics simulations to obtain reliable atomistic ensembles of dynamic protein-RNA complexes.

## MATERIAL AND METHODS

### Protein expression and purification

The Pum-HD and Brat-NHL constructs used in this study comprise residues 1093 to 1426 and 756 to 1037 of the respective proteins (Figure 1B, C), which are connected by a short linker containing a Usp2cc cleavage site to a His_6_-Ubiquitin tag in pHUE plasmid (25,26). The expression was done as previously described (23), but *E. coli* BL-21(DE3) Rosetta cells were used. The cells were then lysed first by incubation for 20 minutes on ice in lysis buffer (50 mM Tris, 1 M NaCl, 5% glycerol, 10 mM imidazole, pH 8.0) with 1 mg/ml lysozyme, 1 μg/ml DNase I, 2 μg/ml Rnase A and protease inhibitor tablets (Roche)), and then by sonication at 4 °C. The lysate was then cleared by centrifugation (18000 g, 4 °C, 1 hour) and purified using a HisTrap HP 5 ml column (GE Healthcare) and a 10-100 mM imidazole linear gradient followed by elution with 250 mM imidazole, which were created by mixing the lysis buffer with 50 mM Tris, 1 M NaCl, 5% glycerol, 1 M imidazole, pH 8.0. Pooled fractions of the proteins were then cleaved by His_6_-tagged Usp2cc at 4 °C overnight during dialysis in dialysis buffer (50 mM Tris, 150 mM NaCl, 1 mM DTT, pH 7.4). Pum-HD was then separated from the tag by reverse affinity chromatography and in the final step Pum-HD was further purified by size-exclusion chromatography (SEC) using HiLoad 16/600 Superdex 75 column (GE Healthcare) in 50 mM Tris, 150 mM NaCl, 1 mM DTT, pH 7.4 buffer. For Brat-NHL serial reverse affinity and heparin chromatography (HiTrap HP Heparin 5ml column – GE Healthcare) followed the affinity chromatography. The sample was loaded on the columns connected in series after equilibrating the columns in dialysis buffer. Brat-NHL was eluted using a 150 mM to 1.5 M NaCl gradient. The Nanos-ZnF plasmid used in this study contains residues 301-392 of Nanos (Figure 1B) connected via a linker with SenP2 cleavage site to a His_6_-SUMO tag in pETM11-SUMO3GFP vector (EMBL Protein Expression and Purification Core Facility). Nanos-ZnF was expressed by transforming the plasmid to *E. coli* BL-21(DE3) Rosetta cells, growing the cells at 37 °C while shaking until OD_600_ of 0.6-1.0, followed by addition of IPTG to 0.3 mM final concentration and further overnight incubation at 16 °C while shaking. Nanos-ZnF was then purified as described above for Brat-NHL, except that in the first affinity chromatography step the basic lysis buffer was 50 mM Tris, 500 mM NaCl, 50 μM ZnSO_4_, 10 mM imidazole, pH 8.0 and the tag was cleaved off using His_6_-tagged SenP2.

### Isotopic labelling

To obtain diversely isotopically labelled proteins, the cells were grown in M9 minimal media and the expression and purification followed the standard protocols outlined above unless stated otherwise. In order to obtain ^15^N-labelled Nanos-ZnF and Brat-NHL, the expression was done using ^15^NH_4_Cl as the sole nitrogen source. In order to obtain various degrees of ^2^H-labelling, cells were grown as follows: a 5 ml overnight culture of H_2_O M9 minimal media with adequate isotopes was spun down, resuspended in 100 ml of D_2_O M9 minimal media with adequate isotopes, grown to an OD_600_ of ∼0.6 at 37 °C while shaking, then diluted to 500 ml final volume with D_2_O M9 minimal medium. The perdeuterated Brat-NHL and Pum-HD for SANS measurements were obtained by expressing the proteins in D_2_O M9 minimal media with ^2^H-glucose as the sole carbon source. For NMR relaxation and ^1^H,^13^C HMQC experiments, Brat-NHL was expressed either uniformly ^2^H-, ^13^C- and ^15^N-labelled (for free form and RNA-bound measurements) or uniformly ^2^H- and ^15^N-labelled (in the *hb* complex) and in both cases with C_δ1_ of isoleucines and one of the methyl groups of valines and leucines ^1^H,^13^C labelled, and the other methyl groups ^2^H-, ^12^C-labelled. The uniform labelling was achieved by expression in D_2_O M9 minimal media with ^15^NH_4_Cl as the sole nitrogen source, and ^2^H- or ^2^H,^13^C-glucose as the sole carbon souce, and adding 2-keto-3,3-d_2_-1,2,3,4-^13^C-butyrate and 2-keto-3-methyl-d_3_-3-d_1_-1,2,3,4-^13^C-butyrate one hour prior induction as described previously (27).

### *Hb* complex formation

To form the *hb* complex, the purified Pum-HD, Brat-NHL and Nanos-ZnF proteins were incubated with a 23 nucleotide-long NRE2 RNA, encompassing the NRE2 in the 3’ UTR of *hunchback* mRNA of the sequence 5’ UUGUUGUCGAAAAUUGUACAUAA 3’ (Microsynth, Figure 1A). The complex was formed by incubating NRE2 RNA with Pum-HD, Nanos-ZnF and Brat-NHL in a 1:1:1:2 molar ratio, respectively, to account for the lower affinity of Brat-NHL towards the RNA. Pum-HD, Nanos-ZnF and NRE2 RNA were diluted to 10 μM, whereas Brat-NHL was diluted to 20 μM. Pum-HD and Nanos-ZnF were then mixed with the RNA, incubated on ice for a 15 minutes prior to addition of Brat-NHL and incubation overnight at 4 °C. The mixture was then concentrated by reducing the volume from 48 ml to 1 ml using a 3 kDa cutoff concentrator and the *hb* complex was purified by SEC using a HiLoad 16/600 Superdex 200 pg column (GE Healthcare). The identity of the *hb* complex peak was confirmed by SDS-PAGE, UV absorption measurement at 260 nm and size-exclusion chromatography-coupled multiangle laser light scattering (SEC-MALLS).

### Isothermal Titration Calorimetry

All isothermal titration calorimetry (ITC) measurements were done on a MicroCal PEAQ-ITC instrument (Malvern) at 20 °C in 50 mM Tris, 150 mM NaCl, 0.5 mM TCEP, pH 7.4 buffer. Diluted NRE2 RNA was snap-cooled before the measurements by incubating at 65 °C, shaking for 5 minutes and quickly cooling on ice for 20 minutes. The proteins were then concentrated and both the protein and RNA solution were degassed. The diluted RNA solution was then added to the cell and the concentrated protein solution was titrated to the RNA from the syringe. Each titration comprised either 13 or 16 injections always with an 0.4 μl initial injection followed by either 2.5 or 3 μl injections. The number of injections was selected to optimize the signal with respect to enthalpy change. The sample was stirred at 750 rpm, instrument feedback was set to high, the reference power was set to 10 μcal/s and the delays were set to 60 second initial delay followed by 150 second delays. The specifics about injections and concentrations in individual titrations are listed in Supplementary Table S1.

### Small-angle scattering

Small-angle X-ray scattering (SAXS) data were acquired at the P12 beamline at DESY, Hamburg, Germany using SEC coupled online to the beamline and a MALLS instrument in parallel. The measurement was done in a 50 mM Tris, 150 mM NaCl, 1 mM DTT, 3% glycerol, pH 7.4 buffer at 25°C. An Agilent BioInert HLPC/FPLC system equipped with Superdex 200 Increase 10/300 GL column (GE Healthcare) was used for the SEC. The *hb* complex at 7.2 mg/ml concentration was injected and run at 0.6 ml/min flowrate. SAXS measurements were then done over a 1 second exposure period per frame using X-rays with 1.23 Å wavelength and Pilatus 6M at 3 m detector distance. The scattering curve was then obtained by averaging frames of the SEC peak with a consistent radius of gyration (*R*_*g*_) and subtracting averaged buffer signal from frames in a baseline region of the SEC run before and after the *hb* complex peak. The MALLS measurements were done on a Wyatt system consisting of miniDAWN TREOS MALLS module with 3 detection angles, Wyatt QELS+ DLS module and Optilab T-rEX refracting index detector operating at 658 nm. The run was analysed using the Zimm fit method in the ASTRA software (Wyatt Technology).

For small-angle neutron scattering (SANS) measurements, samples of the *hb* complex with varying subunit-selective perdeuteration of the protein components were measured at concentrations ranging from 3.7 to 5 mg/ml in 200 μl of 50 mM Tris, 150 mM NaCl, 1 mM DTT, pH 7.4 buffer in Hellma® 100QS quartz cuvette at 20 °C. The samples were generally measured at three different percentages of D_2_O content in the buffer – at 0% D_2_O, at D_2_O percentage close to ^1^H protein match point (∼42%) and at D_2_O percentage close to the ^1^H RNA match point (∼63%). Match points of each sample were calculated using SASSIE-web (https://sassie-web.chem.utk.edu/sassie2/).

The following samples were measured as standards: a boron/cadmium sample for 5 minutes, an empty cell for 10 minutes, and 0%, 42% and 63% D_2_O buffers each for 20 minutes. The samples of the *hb* complex were than measured for 60 minutes each. Transmission of each sample was measured to precisely determine the D_2_O content of the buffer. The measurements were done at the D22 instrument at ILL, Grenoble, France using a neutron wavelength of 6 Å, laterally shifted detector and a detector and collimator distance of 4 m, respectively. The samples are listed in detail in Table 1. The identifier of the experimental dataset is https://doi.ill.fr/10.5291/ILL-DATA.BAG-8-36.

**Table 1.**
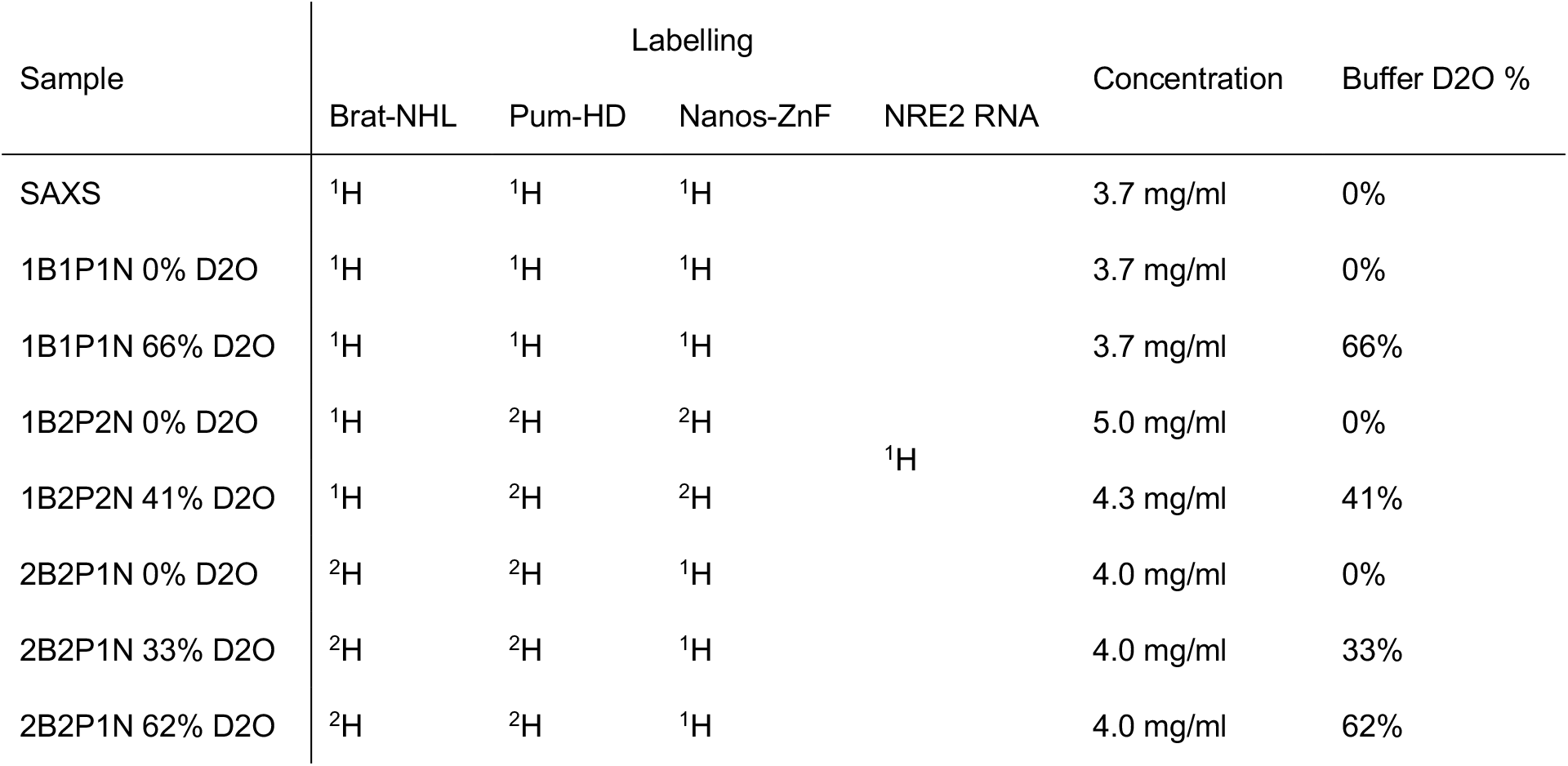
List of samples for small-angle scattering measurements.

All experimental scattering curves were buffer subtracted and initial points with beamstop shadow were removed. The data was processed and analysed using various software from the ATSAS package (28), specifically PRIMUS (29), CRYSON (30), CRYSOL (31) or EOM (32) as well as using ScÅtter (33).

### Cross-linking/mass-spectrometry

Cross-linking of the *hb* complex was done by adapting a previously published protocol (34). The complex was cross-linked by a mixture of ^1^H- and ^2^H-labelled disuccinimidyl suberate (DSS) in 20 mM HEPES, 150 mM NaCl, 1 mM DTT, pH 7.4 buffer. First, 1.5 mg/ml *hb* complex was cross-linked by 0.2 and 1 mM DSS. The cross-linking was done once using 0.2 mM DSS and in three biological replicates using 1 mM DSS. The reaction was incubated while shaking at 37 °C for 30 minutes and then quenched by addition of NH_4_HCO_3_ to 50 mM final concentration. The cross-linked complex was then digested by incubating with LysC (Wako) at 1:100 protease:protein ratio at 37 °C for 3.5 hours and with Trypsin at 1:50 protease:protein ratio at 37 °C overnight. The digested peptides were then desalted using OASIS® HLB µElution Plate and enriched by SEC in 30% (v/v) acetonitrile (ACN) and 0.1% (v/v) trifluoro acetic acid (TFA) using a Superdex Peptide PC 3.2/30 column (GE Healthcare). After evaporating to dryness and dissolving in 4% (v/v) ACN in 1% (v/v) formic acid (FA) the samples were analysed by liquid chromatography-coupled tandem mass spectrometry (LC-MS/MS) using a nanoAcquity UPLC system (Waters) connected online to an LTQ-Orbitrap Velos Pro instrument (Thermo). LC was done using a BEH300 C18 nanoAcquity UPLC column (Waters) with a 3% to 85% (v/v) gradient of ACN in 0.1% (v/v) FA. The MS/MS was done using a top-20 strategy – by first acquiring survey MS scans in m/z range of 375-1600 m/z in the Orbitrap (resolution of 30,000) and fragmenting the top 20 of the most abundant ions per full scan by collision-induced dissociation (CID, with 40% normalized collision energy) and analysing them in the LTQ. Charge states 1, 2 and unknown were rejected to focus the acquisition on larger cross-linked peptides and dynamic exclusion was set to 60 s. Ion target values for full scans and for MS/MS scans were 1,000,000 (for 500 ms max fill time) and 10,000 (for 50 ms max fill time), respectively. The LC-MS/MS was done in two technical duplicates. Since 0.2 and 1 mM DSS cross-linking of 1.5 mg/ml complex produced a high number of multimeric intradomain cross-links, so eventually the cross-linking was repeated in two biological replicates using 0.95 mg/ml *hb* complex and 0.5 mM DSS. The cross-linking, digestion and peptide separation was done as described above, but the mass spectrometry analysis was done using UltiMate™ 3000 RSLCnano system (Thermo Fisher Scientific) directly coupled to an Orbitrap Fusion Lumos (Thermo Fisher Scientific). Dried peptides were dissolved in 4% (v/v) ACN in 1% (v/v) FA and then LC was done using µ-Precolumn C18 PepMap 100 trapping cartridge and nanoEase MZ HSS T3 column in 0.05% (v/v) TFA with a 2% to 85% (v/v) gradient of ACN in 0.1% (v/v) FA. Full scans were acquired with an m/z range of 375-1,600 m/z at 120,000 resolution. For peptide fragment spectra, the quadrupole window was set to 0.8 m/z and the peptides were fragmented by CID (35% normalized collision energy). Charge states 3-7 were selected and dynamic exclusion was set to 60 s. Ion target values for full scans and for MS/MS scans were 200,000 (for 250 ms max fill time) and 20,000 (for 100 ms max fill time), respectively. The spectra were assigned using xQuest and the posterior probabilities were calculated using xProphet (34). The results were then filtered using the following parameters: FDR of 0.05, min delta score of 0.95, −4 to 7 ppm tolerance window and id-score higher than 25. The two biological replicates of 0.95 mg/ml *hb* complex cross-linked at 0.5 mM DSS yielded 20 unique interdomain cross-links.

### NMR spectroscopy

All of the NMR experiments were measured on a Bruker Avance III NMR spectrometer operating at a magnetic field strength corresponding to an 800 MHz proton Larmor frequency, equipped with a Bruker TXI cryo-probe head. The measurements were done at 25 °C in 50 mM Tris, 150 mM NaCl, 1 mM DTT, pH 7.4 buffer. All experiments except Brat-NHL backbone assignment experiments were recorded using apodization weighted sampling (35). Backbone resonance assignment of ^2^H-, ^15^N-, ^13^C-labelled Brat-NHL was achieved to a completion of 77% (excluding prolines) using TROSY-based ^1^H,^15^N-HSQC, HNCA, CBCA(CO)NH and HNCACB triple resonance experiments (36-38). NMR interaction studies were performed as follows: 260 µM ^15^N Nanos-ZnF was titrated by unlabelled Brat-NHL to a molar ratio of 1:1.5, with an intermediate step at 1:1 molar ratio. 170 µM ^15^N Brat-NHL was titrated by unlabelled Nanos-ZnF to a 1:1 molar ratio with no intermediate steps. The experiments were monitored by recording a ^1^H,^15^N-HSQC at each step. The methyl-HMQC spectra comparison of Brat-NHL was obtained by recording a ^1^H,^13^C-HMQC. For free Brat-NHL, methyl-HMQC spectra and ^15^N T_1_ and T_2_ relaxation experiments were measured on a 915 μM ^1^H/^2^H-methyl ILV ^2^H,^13^C,^15^N-labelled Brat-NHL sample. The following delays were used for the T_1_: 20, 50, 100, 150, 400, 500, 650, 800, 1000 and 1200 ms, with the 20 and 150 ms delays measured in duplicates. 12.5, 25, 37.5, 50, 75, 125 and 150 ms delays were used for T_2_ experiment and the 25 ms delay was measured twice. For Brat-NHL bound to RNA, the methyl-HMQC spectra were acquired on a 325 μM 1:1 mixure of ^1^H/^2^H-methyl ILV ^2^H,^13^C,^15^N-labelled Brat-NHL with NRE2 RNA. For Brat NHL in the *hb* complex, the methyl-HMQC, ^15^N T_1_ and T_2_ relaxation measurements were measured on a 65 μM *hb* complex reconstituted using ^2^H-labelled Pum-HD, unlabelled Nanos-ZnF, ^1^H/^2^H,^13^C/^12^C-methyl ILV ^2^H,^15^N labelled Brat-NHL and NRE2 RNA. For the relaxation experiments the following delays were used: 20, 150, 400, 800 and 1200 ms, with the 150 ms delay measured in duplicates for the T_1_ experiment and 12.5, 25, 50, 75 and 125 ms, with the 25 ms delay measured in duplicates for the T_2_ experiment.

All spectra were processed using NMRPipe (39) and analysed using CARA (http://cara.nmr.ch), CcpNmr Analysis (40), or Sparky (41). Peak fitting, error estimation and exponential fitting for the relaxation experiments was done using PINT (42,43). The experimental rotational correlation times were calculated according to Equation 1,

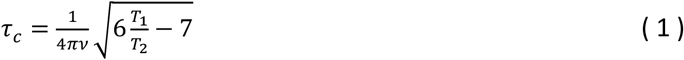

where ν is the Larmor frequency in Hz, *T*_*1*_ is the ^15^N spin-lattice relaxation time and *T*_*2*_ is the ^15^N spin-spin relaxation time (44). The theoretical rotational correlation times were calculated from atomic structures using the ELM module in ROTDIFF 3 (45,46).

### Rigid body modelling

The computational modelling of the *hb* complex was done in Crystallography & NMR System (CNS) (47,48) adapting a previously published protocol (49,50). The starting model of the complex was generated from the published structures of Brat NHL-RNA complex (4zlr) (22) and Pum HD-Nanos ZnF-RNA complex (5kl1) (15) by threading the individual complexes on a single NRE2 RNA containing both binding sites and the connecting nucleotides. A pool of models of the complex was then generated by modified protocols within ARIA (51). First, the conformation of the NRE2 RNA nucleotides, which are not bound in either of the published RNA complexes, was randomized. The conformation of the proteins and the RNA elucidated in the published structures were then kept fixed throughout the modelling. The randomized structures were minimized unrestrained or with XL/MS derived distances employing standard simulated annealing protocols within ARIA. The pool of unrestrained models was then generated by 40,000 MD steps of 6 fs each, whereas the pool of models with XL/MS as distance restraints was generated by 120,000 MD steps of 2 fs each. Unrestrained modelling generated 5,055 models, modelling with XL/MS data generated 4,572 models. The XL/MS data were put in as 30 Å lysine-lysine C_α_-C_α_ upper distance limits with log harmonic potential (52). Each model was then fitted against the experimental data using CRYSON (30) and CRYSOL (31), then for each modelling an ensemble of best fitting models was selected according to a criterion of χ^2^ values of the fits of the model to each experimental curve. In case of unrestrained modelling the criterion 0.23 quantile of the lowest χ^2^ values, whereas for modelling with XL/MS data the criterion used was 0.3 quantile of the lowest χ^2^ values.

### All-atom molecular dynamics simulations

Eight different conformations of the protein-RNA complex were taken from the rigid-body modelling and henceforth used as starting conformations for all-atom explicit-solvent simulations. MD simulations were carried out with the Gromacs software (53), version 2019.6. Interactions of the protein and the RNA were described with the Amber99sbws force-field, and the TIP4P-2005s water model was used (54). Each of the eight initial conformations was placed in a dodecahedral simulation box, where the distance between the protein to the box edges was at least 2.0 nm. The boxes were filled with 184,665 water molecules, and 17 sodium ions where added to neutralize the systems. In total, the simulations systems contained 751,094 atoms. The energy of each simulation system was minimized within 400 steps with the steepest decent algorithm. Subsequently, the systems were equilibrated for 100 ps with harmonic position restraints applied to the backbone atoms of the proteins (force constant 1000 kJ mol^−1^nm^−2^). Finally, each of the eight replicas was simulated for 110 ns without any restraints. The temperature was kept at 298 K using velocity rescaling (*τ* = 0.1 ps) (55). The pressure was controlled at 1 bar with the Berendsen (*τ* = 1 ps) (56) and with the Parrinello-Rahman barostat (*τ* = 5 ps) (57) during equilibration and production simulations, respectively. The geometry of water molecules was constrained with the SETTLE algorithm (58), and LINCS was used to constrain all other bond lengths (59). Hydrogen atoms were modeled as virtual sites, allowing a time step of 4 fs. The Lennard-Jones potentials with a cut-off at 1nm were used to describe dispersive interactions and short-range repulsion. Electrostatic interactions were computed with the smooth particle-mesh Ewald method (60,61). Visual inspection of the simulations revealed that the RNA–protein contacts were stable throughout the simulations.

### Explicit-solvent SAXS/SANS calculations

The SAXS and SANS calculations were performed with an in-house modification of Gromacs 2018.8, as also implemented by our webserver WAXSiS (62-64). The implementation and tutorials are available at https://biophys.uni-saarland.de/software.html. Simulation frames from the time interval between 30 and 110 ns were used for SAXS/SANS calculations. A spatial envelope was built around all solute frames from all eight replicas of the protein-RNA complex. Solvent atoms inside the envelope contributed to the calculated SAXS/SANS curves. The distance between the protein-RNA complex and the envelope surface was at least 1.0 nm, such that all water atoms of the hydration shell were included. The buffer subtraction was carried out using 783 simulations frames of a pure-water simulation box, which was simulated for 110 ns and which was large enough to enclose the envelope. The orientational average was carried out using 1700 **q**-vectors for each absolute value of *q*, and the solvent electron density was corrected to the experimental value of 334 e/nm^3^, as described previously (63). During SANS calculations, the perdeuteration conditions and D_2_O concentrations were taken according to the experimental conditions. Here, we assigned the mean neutron scattering length to all potentially deuterated hydrogen atoms, as described previously (62). This protocol leads to a constant offset in the SANS curves that is absorbed into a fitting parameter (see below). To compare the experimental with the calculated SAXS/SANS curves, we fitted the experimental curve via *I*_exp,fit_(*q*) = *f*·*I*_exp_ + *c*, by minimizing the chi-square with respect to the calculated curve. Here, the factor *f* accounts for the overall scale, and the offset *c* takes the uncertainties from the buffer subtraction and the incoherent scattering in SANS experiments into account. No fitting parameters owing to the hydration layer or excluded solvent were used, implying that also the radius of gyration was not adjusted by the fitting parameters.

## RESULTS

### Brat-NHL and Pum-HD do not bind NRE2 cooperatively

To test if there is cooperativity between Pum-HD and Brat-NHL in RNA binding, we quantified the influence of Pum-HD on the interaction of Brat-NHL with *hunchback* mRNA by ITC. First, we titrated Brat-NHL to a 23-mer NRE2 RNA covering NRE2 of *hunchback* mRNA 3’ UTR and including Brat and Pumilio binding sites (Figure 1A). Brat-NHL binds to this RNA with a K_d_ of 1.2 ± 0.2 μM (Figure 1D). Incubating the RNA with Pum-HD was suggested to increase the affinity of Brat-NHL to the RNA based on electrophoretic mobility shift assays (EMSA) (22). Thus, in the second experiment we incubated the NRE2 RNA with equimolar amounts of Pum-HD and then titrated Brat-NHL to the mixture (Figure 1E). The K_d_ of Brat-NHL to NRE2 RNA in the presence of Pum-HD apparently increased to 2.9 ± 1.4 μM. Due to a larger error it cannot be clearly concluded that the presence of Pum-HD decreases the affinity of Brat-NHL to NRE2 RNA. After taking the margin of the error into account, the results of this experiment at least demonstrate a lack of cooperativity in RNA binding between Brat-NHL and Pum-HD. It was speculated that Pum-HD might increase the affinity of Brat-NHL to the RNA by melting the secondary structure of the RNA. Therefore, it should be noted here that the purified Pum-HD purified by us was positively tested for the ability to unfold the RNA prior to the ITC experiments and binds the NRE2 RNA with high affinity (Supplementary Figure S1, Figure 1F). Moreover, extending an RNA with the Brat-NHL binding site by multiple uracils was shown to increase the affinity of Brat-NHL to the RNA, supposedly due to multivalent binding (22). However, extending the Brat-NHL binding site to NRE2 RNA did not noticeably increase the affinity of Brat-NHL (Figure S2B vs. Figure S2F), confirming that Brat-NHL is not able to use NRE2 RNA as a multivalent platform. Full ITC data including heat profiles are shown in Supplementary Figure S2.

### Brat-NHL does not interact with Nanos-ZnF *in vitro*

As no cooperativity between Brat-NHL and Pum-HD in RNA binding was observed, we then wondered if cooperativity is instead mediated by Nanos-ZnF. Pum-HD and Nanos-ZnF bind the *hunchback* mRNA cooperatively (15), so an interaction between Nanos-ZnF and Brat-NHL could extend this cooperativity to Brat-NHL. Therefore, we tested a direct interaction between Nanos-ZnF and Brat-NHL using NMR titrations. Typically, NMR titrations are applicable to a very broad range of interaction affinities from tight to weak or transient interactions, so we selected this method to avoid missing a potential interaction by probing a limited K_d_ range. We expressed both Brat-NHL and Nanos-ZnF ^15^N-labelled as well as unlabelled and then titrated either ^15^N-labelled protein by their unlabelled counterpart, while monitoring the titration by ^1^H,^15^N HSQC spectra (Figure 2). Neither of the two NMR titrations revealed any signs of interaction. We did not observe any changes of peak positions, that would be indicative of changes of the local chemical environment accompanying an interaction, or changes in peak intensity, reflecting changes of molecular tumbling upon complex formation. It should be noted that the Nanos-ZnF construct is one third of the molecular weight of the Brat-NHL construct (32 kDa), so a 4-fold increase of the apparent molecular weight upon complex formation should change molecular tumbling of Nanos-ZnF to a degree that obvious changes of peak intensities in NMR should be observed.

**Figure 2.**
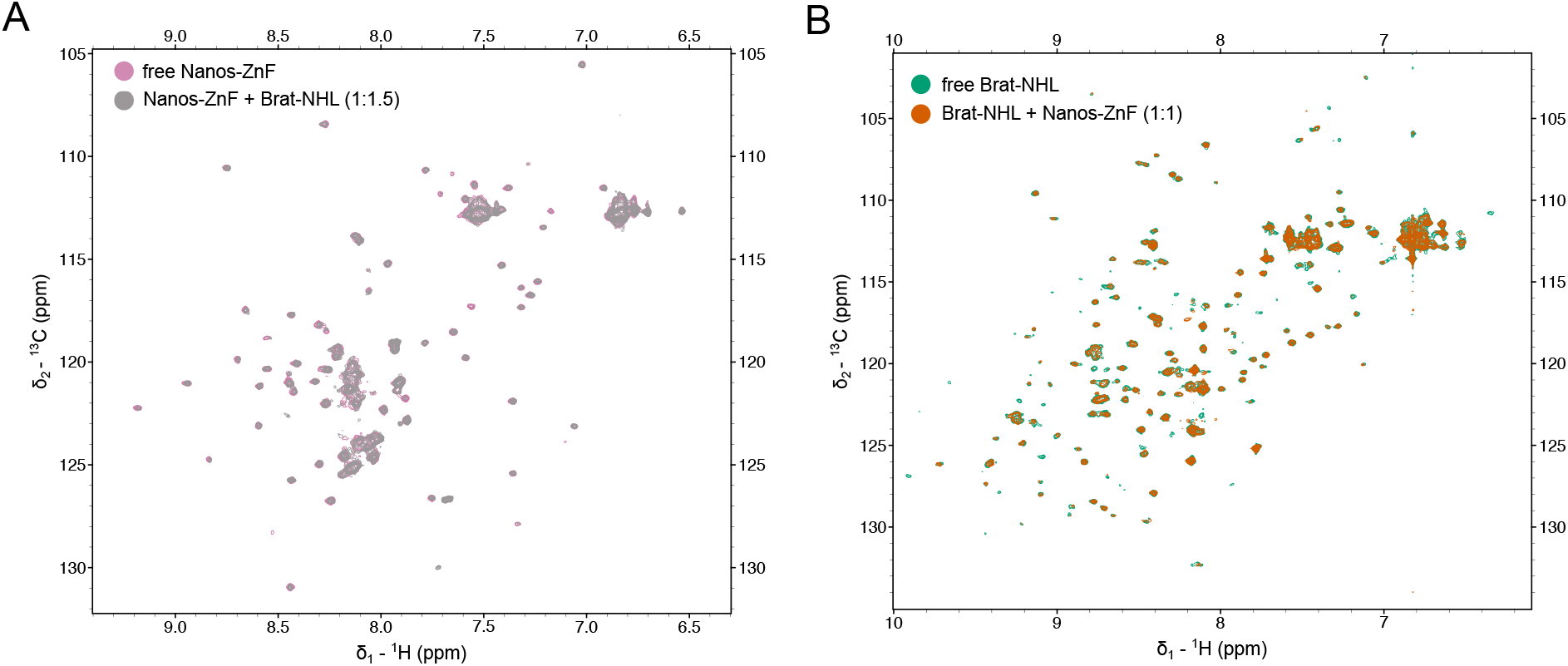
Brat-NHL and Nanos-ZnF do not interact in the absence of RNA. NMR titrations were performed to test if Brat-NHL and Nanos-ZnF interact *in vitro* in the absence of RNA. **(A)** Overlay of the ^1^H,^15^N HSQC spectra of ^15^N labelled Nanos-ZnF in the free form and with unlabelled Brat-NHL added to 1:1.5 Nanos-ZnF:Brat-NHL molar ratio. **(B)** Overlay of the ^1^H,^15^N HSQC spectra of ^15^N labelled Brat-NHL in the free form and with unlabelled Nanos-ZnF added to 1:1 Brat-NHL:Nanos-ZnF molar ratio. The spectrum of the respective free proteins is always rendered with a lower contour limit, so it is visible that two spectra are overlaid and no chemical shift perturbations or signal intensity decrease is observed in either of the experiments.

### Pum-HD, Nanos-ZnF and Brat-NHL assemble on a single NRE2 RNA

Interestingly, when Pum-HD and Nanos-ZnF recognize *hunchback* mRNA cooperatively the structure of the complex reveals an interaction between Pum-HD and Nanos-ZnF that only exist in the presence of RNA (15). Therefore, we tested if Pum-HD, Nanos-ZnF, Brat-NHL and *hunchback* mRNA form a quaternary *hb* complex. NRE2 RNA was incubated with Pum-HD, Nanos-ZnF and Brat-NHL in a 1:1:1:2 ratio, and purified as described above. The expected molecular weight of an equimolar quaternary complex of the NRE2 RNA with the three domains is approximately 87.8 kDa. The SEC chromatogram showed a major peak with a maximum at an elution volume corresponding to approximately 77 kDa. The peak contains all three protein domains, as visible by SDS-PAGE, and the NRE2 RNA, which is evidenced by a strong UV absorption at 260 nm. The peak was further analysed by SEC-MALLS-SAXS to gain further insights into the architecture of the complex. Notably, MALLS reveals a monodisperse particle with a molecular weight of 85.0 ± 1.7 kDa (Supplementary Figure S3). Collectively, these results confirm that the four components form a stable 1:1:1:1 *hb* complex of Pum HD, Nanos ZnF, Brat NHL and NRE2 RNA.

### Structural characterization of the *hb* complex

Next, we set out to investigate the *hb* complex using X-ray crystallography. We tested an extensive set of commercial crystallization screens covering various crystallization chemistry (PEGs, salts, PEG smears, alternative polymer precipitants, etc.) at a wide range of *hb* complex concentrations and multiple temperatures. In addition, we designed and tested custom screens around the published crystallization conditions of Pum-HD, Nanos-ZnF and Brat-NHL (all conditions tested are listed in Supplementary Table S2). The extensive testing did not yield any diffracting crystals of the *hb* complex. Lastly, carrier-driven crystallization using a Pum-HD fused to an MBP-tag also failed, likely because the modification of Pum-HD interfered with complex formation. We then adopted an alternative integrative approach to obtain a structural model of the *hb* complex, which combines small-angle X-ray and neutron scattering (SAXS/SANS), NMR spectroscopy, cross-linking/mass-spectrometry (XL/MS), available crystal structures of Brat-NHL-RNA complex and Pum-HD-Nanos-ZnF-RNA complex (15,22), as well as molecular dynamics simulations.

First, SAXS data indicate a structured particle with an R_g_ of ∼37.4 Å according to Guinier analysis (Supplementary Figure S4A). Indirect Fourier transformation results in an asymmetric distance distribution function (P(r)) with a maximum at ∼30 Å that smears out through an additional peak to a maximum particle dimension (D_max_) of ∼130 Å (Figure 3B). This is smaller than the sum of D_max_ of the components (Supplementary Figure S4B). The P(r) of Brat-NHL-RNA complex is a single approximately symmetrical peak with a maximum at ∼25 Å and a *D*_*max*_ of ∼50 Å. The P(r) of the Pum-HD-Nanos-ZnF-RNA complex shows a more complex distribution with a maximum at ∼22 Å and a long smeared tail with an additional peak and a *D*_*max*_ of ∼90 Å. To learn whether the system possesses some degree of flexibility, the SAXS data were also plotted as a dimensionless Kratky plot (Figure 3C). This plot reveals approximately a Gaussian peak, which is typical for a folded globular biomolecule. However, the position of the maximum of the peak deviates from a maximum expected for an ideal globular folded protein indicating either a deviation from a globular shape or some degree of flexibility. In SANS measurements, subunit-selective perdeuteration combined with varying D_2_O concentration in the buffer is used during contrast matching to determine further structural parameters (e. g. centre-of-mass distances between the subunits) (65), which are useful in data-driven structure modelling of protein-RNA complexes (50,66-68). Here, we used several differentially subunit-selectively perdeuterated *hb* complex samples (Table 1). We obtained in total seven SANS scattering curves on a fully protonated *hb* complex, *hb* complex with perdeuterated Pum-HD and Nanos-ZnF, and *hb* complex with perdeuterated Brat-NHL and Pum-HD (Figure 3A).

**Figure 3.**
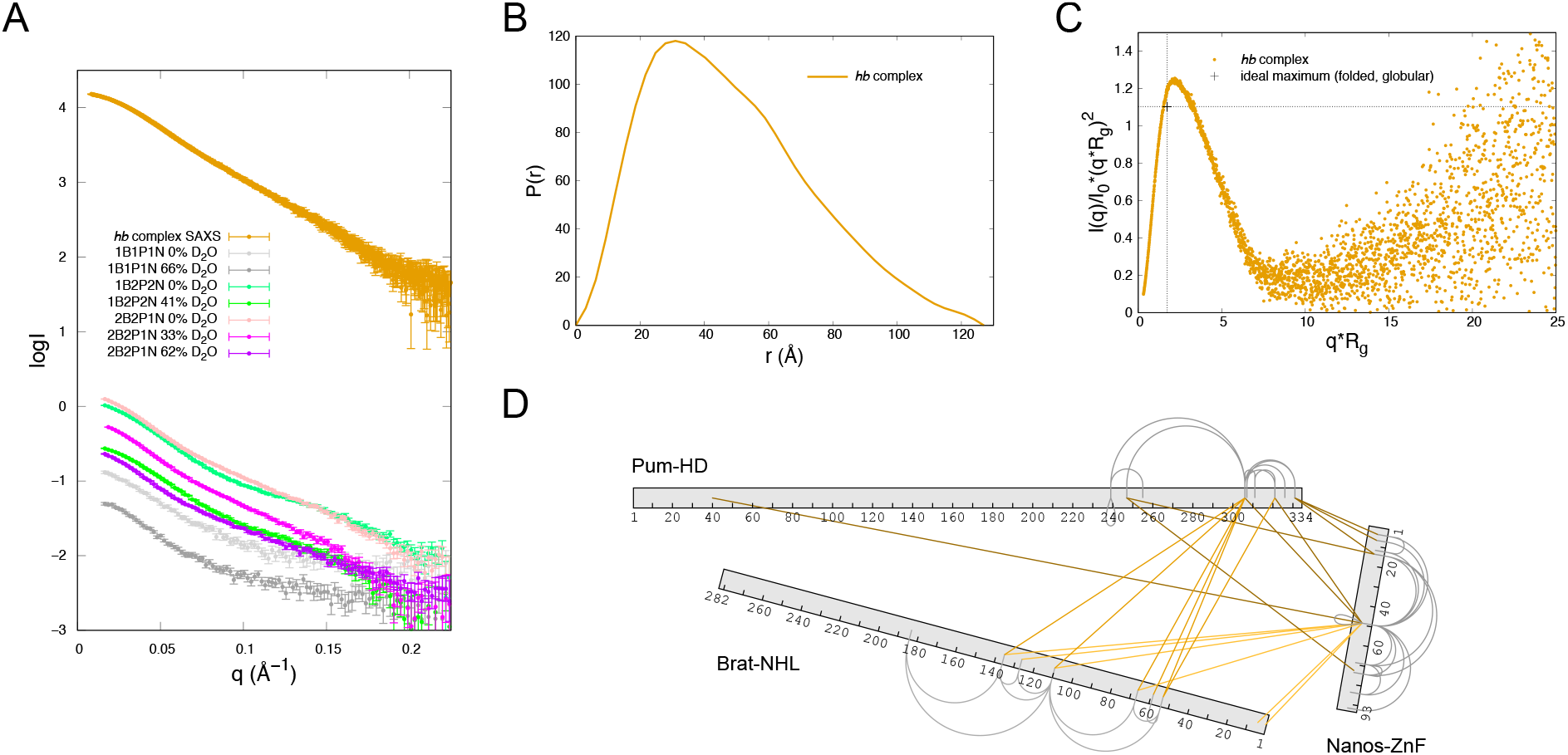
Characterization of the *hb* complex by small-angle scattering (SAS) and cross-linking/mass spectrometry (XL/MS). **(A)** Scattering curves of the *hb* complex. The curve labelled *hb* complex SAXS corresponds to the scattering curve obtained for fully protonated *hb* complex using small-angle X-ray scattering (SAXS) coupled online to SEC. The remaining curves correspond to the scattering curves of *hb* complex obtained using small-angle neutron scattering (SANS). Each curve was measured on a distinct sample of the *hb* complex, that varied in perdeuteration of the components of the complex (Table 1). The perdeuteration is always indicated in the label – 1 stands for a fully protonated component, whereas 2 indicates a perdeuterated complex. B, P and N stands for Brat-NHL, Pum-HD and Nanos-ZnF, respectively. NRE2 RNA was always protonated and all curves shown here were measured at varying concentrations of D_2_O in the buffer as stated in the legend. **(B)** Distance distribution function of the *hb* complex obtained from indirect Fourier transformation of the SAXS scattering curve. **(C)** Normalized Kratky plot of the SAXS scattering curve. The indicated ideal maximum corresponds to an expected maximum for a folded globular protein (86). **(D)** Cross-links of the *hb* complex obtained from the XL/MS experiments. Inter-molecular cross-links are indicated by lines in shades of oranges. Grey loops indicate intra-molecular cross-links. Only cross-links with an id-score higher than 25 are shown.

To search for a potential structure of the *hb* complex described by the experimental SAXS/SANS data, we generated a pool of ∼5000 models by unrestrained molecular dynamics (MD) in CNS using NRE2 RNA, the structure of Brat NHL-RNA complex (PDB ID: 4zlr) (22) and Pum HD-Nanos ZnF-RNA complex (PDB ID: 5kl1) (15). During structure calculation the unbound NRE2 RNA nucleotides (7-11) were randomized and the known structures preserved, including protein-RNA contacts. The pool of structures covers a large conformational space, which we then aimed to reduce comparing the back-calculated scattering curve of each model with the experimental data using CRYSOL and CRYSON (30,31). For each experimental curve we set a cut-off criterion of 0.23 quantile of lowest χ^2^ values of the fit between the experimental and back-calculated scattering, and then searched for models that fulfil the criterion for each experimental curve (SAXS and SANS). This yielded an ensemble of fifteen models, which fall into two clusters representing two alternative conformations, that seem to fit the data equally well (Figure 4A). These two alternative conformations were present even when a stricter cut-off criterion was chosen. Two distinct conformations fitting the data equally well could be a consequence of either the general low resolution of the method or conformational heterogeneity of the complex. The low resolution would allow two distinct conformations to fit well, if the shapes of the conformations are roughly symmetric (this is an intrinsic limitation of the methods and has been extensively described elsewhere (69-71)). In case of conformational heterogeneity, the measured scattering curve is a linear sum of the scattering of individual conformations weighted by their population. In case of equally populated conformations, fitting each conformation individually to the experimental data would result in equally good fits. To resolve this, we acquired more data using XL/MS.

**Figure 4.**
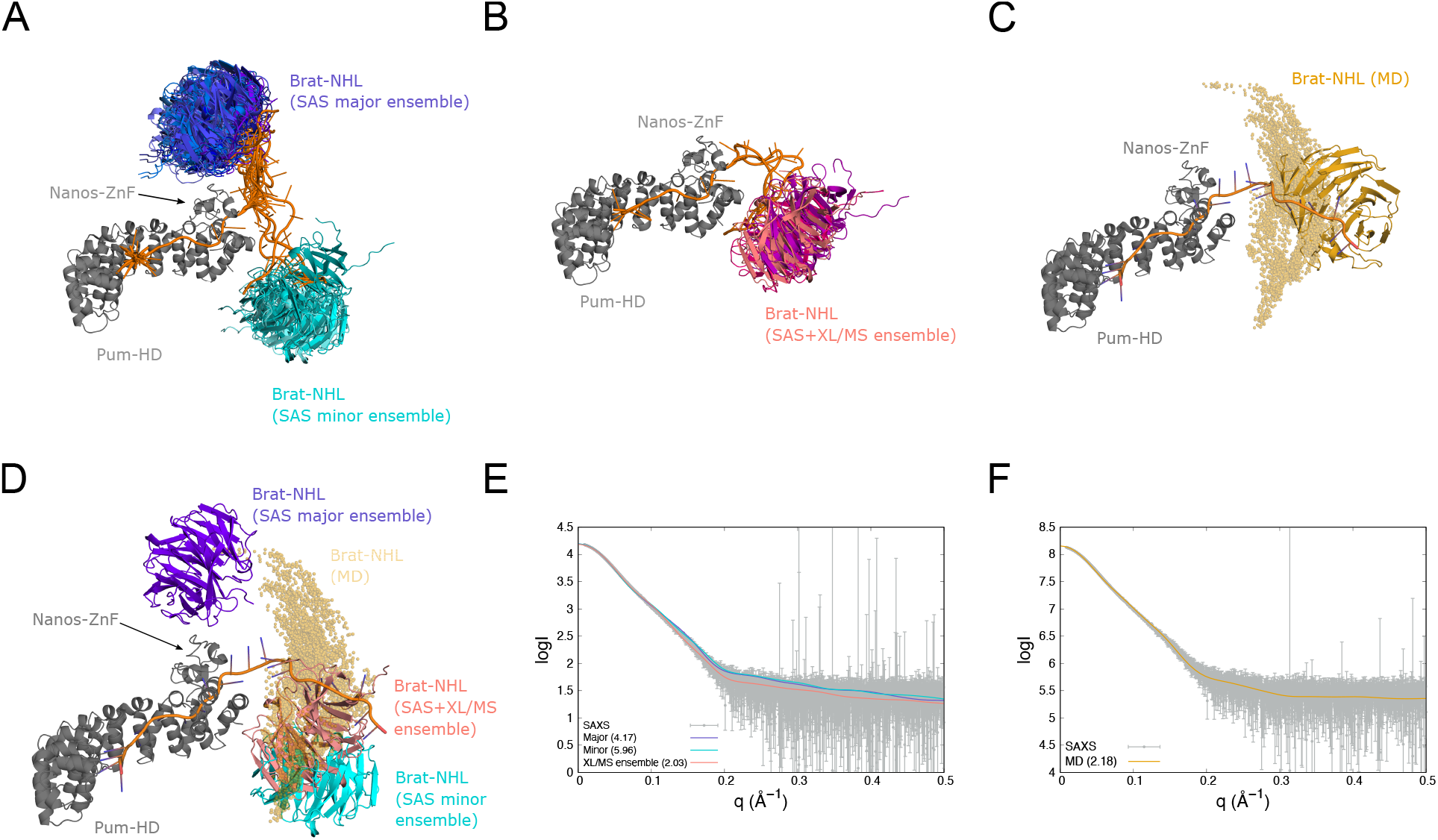
Modelling of the *hb* complex. **(A)** Ensemble of *hb* complex models fitting the SAS data best. The models fall into two clusters representing two distinct conformations. The first, major cluster is shown with Brat-NHL in shades of purple and contains eleven models with Brat-NHL close to the C-terminus of Nanos-ZnF. The second, minor cluster is shown with Brat-NHL in shades of cyan and includes four models with Brat-NHL close to the N-terminus of Nanos-ZnF. **(B)** Ensemble of *hb* complex models derived using a combination of the SAS and XL/MS data. The ensemble comprises three models all adopting the same conformation with Brat-NHL close to the N-terminus of Nanos-ZnF. Individual models are shown with Brat-NHL in shades of magenta. **(C)** Ensemble of *hb* complex models obtained by all-atom MD simulations. One conformation of Brat-NHL is shown in yellow cartoon for illustration, while Brat-NHL domains from each other frame are shown as yellow spheres representing the centre-of-mass of the domain. **(D)** Overlay of *hb* complex models. Representative models of ensembles in (A) and (B) are displayed and overlayed on the ensemble from (C). **(E)** Fits of back-calculated scattering curves of representative models of *hb* complex ensemble to the experimental SAXS curve. ‘Minor’ and ‘Major’ denotes representative models of the two clusters in (A) and ‘XL/MS ensemble’ denotes the representative model of the ensemble in (B). The representatives are the models closest to the mean structure of the ensemble. **(F)** Comparison of SAXS curve computed from all-atom MD with the experimental SAXS curve. All models in the ensembles are always superimposed on Pum-HD and Nanos-ZnF. Numbers in the brackets are the reduced χ^2^ values of the fit.

The cross-linking reaction was optimized to 0.5 mM DSS on 0.95 mg/ml *hb* complex, which yielded twenty inter-molecular cross-links with an id-score higher than 25. Eight of the cross-links were between Pum-HD and Nanos-ZnF, leaving twelve cross-links between Brat-NHL and Pum-HD or Nanos-ZnF. These were then used as lysine-lysine distance restraints with a log-harmonic potential in the same MD protocol as described above to generate another pool of ∼4500 models. The pool of models was then reduced as above, but with a cut-off criterion of 0.3 quantile of lowest χ^2^ for each curve (Supplementary Figure S5C, D). Finally, an ensemble was obtained from the restricted pool by selecting only the models in 0.1 quantile of the lowest energy of XL/MS distance restraints violations. The ensemble comprised three models in a conformation similar to the conformation of the minor cluster described above, except that Brat-NHL was rotated by about 45° (Figure 4D). However, even the model with the least violations of XL/MS distance restraints satisfies only about six of the twelve cross-links (Supplementary Figure S5E) and visual inspection of other models revealed that some cross-links can only be satisfied exclusively. Such results would be expected for a flexible system, where there is no fixed position of Brat-NHL relative to the Pum-HD-Nanos-ZnF moiety.

We then used NMR spectroscopy in order to probe the flexibility of the *hb* complex experimentally and quantitatively at residue resolution. Due to the size of the *hb* complex, we used a sample of the complex containing unlabelled Nanos-ZnF and RNA, ^2^H-labelled Pum-HD and ^1^H,^13^C-methyl-ILV, ^2^H, ^15^N-labelled Brat-NHL (Brat-NHL in the complex), and compared NMR measurements with data from samples of free and RNA-bound ^1^H,^13^C-methyl-ILV, ^2^H,^13^C,^15^N-labelled Brat-NHL. These labelling schemes were necessary to compensate signal loss due to size-dependent line broadening. All samples would only provide signal from Brat-NHL, but would report on the dynamic properties of Brat-NHL in its free form, bound to RNA and in the *hb* complex. Two sets of experiments were recorded on the samples – ^1^H,^15^N T_1_ and T_2_ relaxation experiments, and ^1^H,^13^C-HMQC spectra. The relaxation experiments were acquired on free Brat-NHL and on Brat-NHL in the complex. Brat NHL is a disc-shaped ∼32 kDa β-propeller domain, whereas the measured molecular weight of the *hb* complex is ∼85 kDa. Thus, forming the *hb* complex should be accompanied by a drastic change of relaxation properties of Brat-NHL, if the complex tumbles as one entity. However, the acquired relaxation parameters reveal that neither R_1_ (1/T_1)_ nor R_2_ (1/T_2_) changes drastically between free Brat-NHL and Brat-NHL in the complex (Figure 5A and C). We used these values to calculate the rotational correlation time (τ_c_), which is the time it takes the average molecule to rotate one radian in solution, and so it reports on the overall tumbling of the molecule and allows apparent molecular weight estimation (assuming isotropic tumbling). Brat-NHL has a theoretically predicted τ_c_ of 15.3 ns, whereas the predicted τ_c_ of the *hb* complex is 54 ns (with a standard deviation of 1.5 ns, assuming the whole complex tumbles jointly). Both values were predicted using ROTDIFF 3 (45,46)). The τ_c_ of the *hb* complex was calculated as a median value of the seventeen models generated by modelling using only the small-angle scattering data (described above). The experimentally measured τ_c_ of free Brat-NHL of 19.0 ± 4.3 ns matches the predicted value well. Strikingly, the measured τ_c_ of Brat-NHL in the complex was 15.6 ± 3.8 ns, which is similar to the value of free Brat-NHL and is far below the predicted value for a rigid *hb* complex (Figure 5B). The elevated τ_c_ of free Brat-NHL compared to Brat-NHL in the complex may result from weak unspecific multimerization of free Brat-NHL. Nevertheless, these τ_c_ values confirm that the *hb* complex is largely flexible and that Brat-NHL tumbles independently of the Pum-HD-Nanos-ZnF moiety. In such a scenario the measured small-angle scattering curve would be an average over all conformations and the cross-links would arise from stochastic encounters of the domains possibly yielding conflicting data. Additionally, we would not expect any further cooperativity or interaction upon recruitment of Brat-NHL to the complex. To test this, we performed two further experiments. First, we reconstituted the previously characterized Pum-HD-Nanos-ZnF-RNA complex on NRE2 RNA and titrated Brat-NHL to it using ITC (Figure 6A). The K_d_ was 1.4 ± 0.2 μM, which is almost the same to the K_d_ measured for the interaction of Brat-NHL with only NRE2 RNA (1.2 ± 0.2 μM), demonstrating a lack of cooperativity. Secondly, we measured and compared the methyl region of the ^1^H,^13^C TROSY-HMQC spectra of free Brat-NHL, NRE2 RNA-bound Brat-NHL and Brat-NHL in the complex (Figure 6B,C). A comparison of the spectra of free and RNA-bound Brat-NHL shows obvious chemical shift perturbations (CSP). However, a comparison of the spectra of RNA-bound Brat-NHL and Brat-NHL in the complex reveals no further CSP, suggesting that Brat-NHL does not form any additional interaction surface in the *hb* complex with another protein or RNA component.

**Figure 5.**
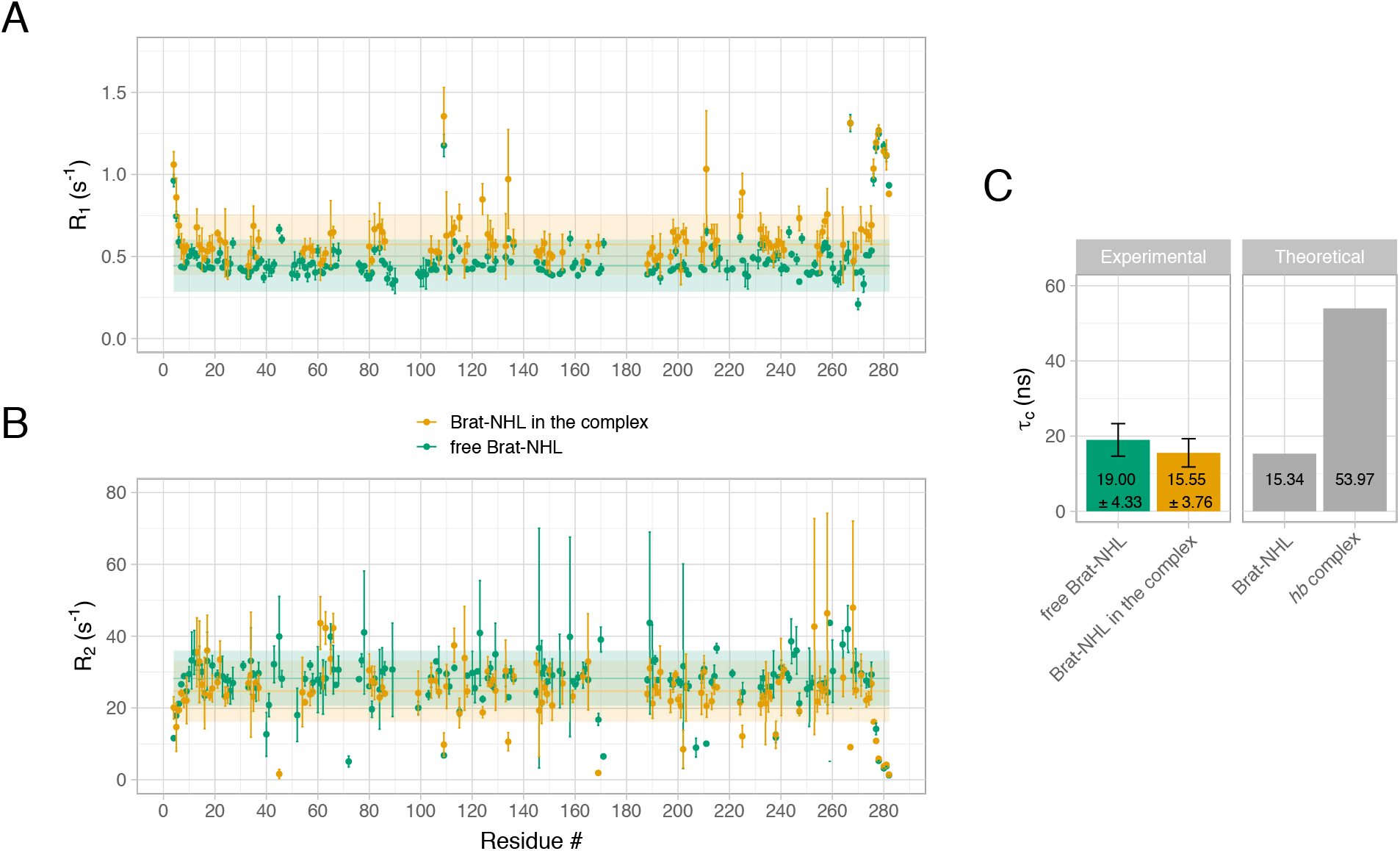
Flexibility of the *hb* complex. **(A)** ^15^N Spin-lattice relaxation rate (R_1_) of Brat-NHL. R_1_ was measured for both Brat-NHL in its free form (Free Brat-NHL) and in the *hb* complex with all the other components of the complex unlabelled (Brat-NHL in the complex). Values related to free Brat-NHL are displayed in green, whereas values related to Brat-NHL in the complex are displayed in orange. Measured values are plotted as dots with error bars, horizontal full line shows a median and the shaded horizontal bar indicates a range of two standard deviations of the median. **(B)** ^15^N Spin-spin relaxation rates (R_2_) of Brat-NHL. R_2_ was measured on the same samples as R_1_, and labelling corresponds to (A). **(C)** Rotational correlation time (τ_c_) of Brat-NHL. The left panel of the plot indicates experimentally determined τ_c_ values calculated based on measured R_1_ and R_2_ values. The right panel shows theoretically expected τ_c_ values calculated from the crystal structure of Brat-NHL or an average of theoretically expected τ_c_ calculated for the ensemble of fifteen models of the *hb* complex modelled using the small-angle scattering data (Figure 4A).

**Figure 6.**
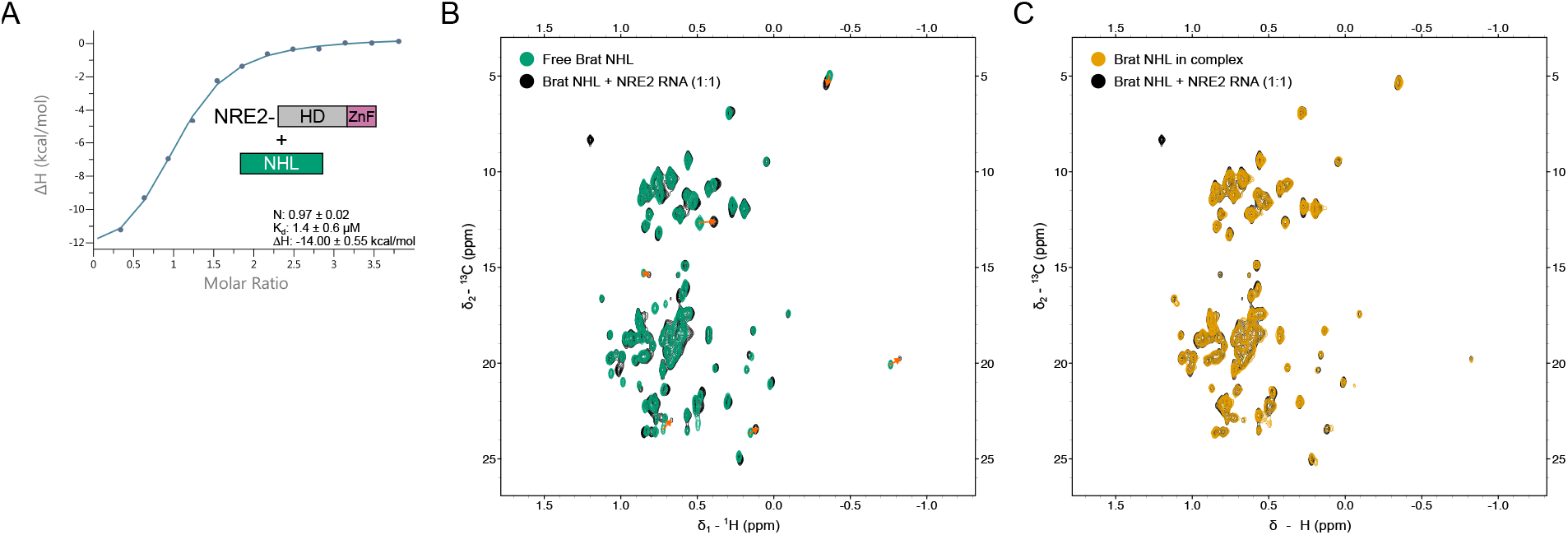
Brat-NHL does not cooperatively interact with Pum-HD and Nanos-ZnF within the *hb* complex. **(A)** Interaction of Brat-NHL with the Pum-HD-Nanos-ZnF-NRE2 RNA complex by isothermal titration calorimetry (ITC) measurements. To test cooperativity of Brat-NHL with Pum-HD and Nanos-ZnF in RNA binding the Pum-HD-Nanos-ZnF-NRE2 RNA complex was formed as previously described (15) and purified by size-exclusion chromatography. Brat-NHL was then titrated to it in an ITC measurement, which revealed an interaction with a K_d_ of 1.4 ± 0.6 μM. The K_d_ of the interaction between free NRE2 RNA and Brat-NHL was 1.2 ± 0.2 μM (Figure 1). **(B)** Overlay of the methyl region of ^1^H,^13^C HMQC spectra of Brat-NHL in its free form (Free Brat-NHL) and Brat-NHL with NRE2 RNA (at 1:1 stochiometric ratio). Red arrows are used to illustrate some of the chemical shift perturbations. **(C)** Overlay of the methyl region of ^1^H,^13^C HMQC spectra of Brat-NHL with NRE2 RNA (at 1:1 stochiometric ratio) and of Brat-NHL in the *hb* complex (Brat-NHL in the complex). No chemical shift perturbations are seen between these two states indicating that no further contacts are formed between Brat-NHL and Pum-HD-Nanos-ZnF.

To test whether a continuous structural ensemble is compatible with the SAXS/SANS data, and to obtain an atomistic model of the overall complex in solution, we used all-atom MD simulations with the Amber99sbws/TIP4P-2005s force field, which has been refined to balance protein-protein versus protein-water interactions (54). To improve the sampling of the conformational space, we simulated 8 replicas for 110ns, where each replica started from a different conformation taken from the rigid-body simulations. No bias or restraint was applied. In the simulations, the conformational space adopted by Brat-NHL relative to Pum-HD-Nanos-ZnF is characterized by an arch-shaped distribution, in which Brat-NHL (i) forms only occasional contacts with Pum-HD-Nanos-ZnF and (ii) takes various rotational states relative to Pum-HD (Figure 4C). SAXS and SANS curves were computed from a total of 6408 simulation frames using explicit-solvent calculations, taking atomistic representations for the hydration layer and excluded solvent into account. The experimental curves *I*_exp_(*q*) were fitted to the calculated curve via *I*_exp,fit_(*q*) = *f*· *I*_exp_(*q*)+*c*, but no parameters related to the hydration layer or excluded solvent were adjusted. Remarkably, we found nearly quantitative agreement between the calculated and experimental SAXS curves (reduced χ^2^ = 2.18) and reasonable agreement for the SANS curves (Figure 4F and Supplementary Figure S5F), without the need of reweighting the ensemble or otherwise coupling the simulation to the data, highlighting the quality of the applied force field and suggesting that the accessible conformational space was reasonably well sampled. For comparison, we also used the established ensemble optimization method (EOM) (32). EOM selected an ensemble of 7 models (Supplementary Figure S6) using the SAXS curve and the pool of random models generated previously for the modelling using only SAS data. The EOM ensemble has an R_flex_ of 88.3 % (R_flex_ of the pool: 89.5 %) and R_σ_ of 5.26, which shows that restricting a pool of approximately 5000 random generated models to an ensemble of 7 models with the use of experimental SAXS data does not reduce the represented conformational space at all. These results suggest that the *hb* complex adopts a continuous, heterogenous ensemble in solution.

Taken all together our data clearly show that a stable quaternary 1:1:1:1 complex of Pum-HD, Nanos-ZnF, Brat-NHL and NRE2 RNA forms. However, the RNA binding of Brat-NHL is completely independent of Pum-HD and Nanos-ZnF and no additional interaction between Brat-NHL and Pum-HD-Nanos-ZnF moiety is observed. The resulting *hb* complex is flexible and the flexibility is presumably only limited by the sterical restrictions of the conformation of the unbound nucleotides between the respective protein binding sites in NRE2 RNA.

## DISCUSSION

Protein-RNA complexes play key roles at any stage of gene expression and underlying molecular mechanisms of their function have been elucidated for large machines like ribosomes, RNA polymerases or the spliceosome (72-74). However, more than 1000 proteins were identified to bind mRNA and influence its fate (75,76). In these cases our understanding of molecular mechanisms of the function of those proteins is mostly limited to insights from structures of individual RNA binding proteins (RBPs) bound to a short cognate RNA (77), but those RBPs do not function in isolation. Only a few studies showed examples of cooperative RNA-recognition by multiple RBPs resulting in increased affinity and specificity (78,79). However, other mechanisms may be possible and detailed mechanistic understanding of complexes beyond those of individual RBPs is essential to formulate general molecular mechanisms governing the mRNA interactome. Shedding light into simultaneous RNA-recognition by multiple RBPs is easily hurdled by the transient and dynamic nature of those complexes. Addressing this challenge often requires an integrative approach combining multiple methods (80). In this study we used such an approach to obtain results allowing us to provide proper interpretation of previously published data and ultimately clarify our understanding of the structure and dynamics of the quaternary protein-RNA complex that suppresses *hunchback* mRNA translation.

In case of Pumilio and Nanos there is now sufficient evidence to conclude that these two proteins indeed act jointly (15), but how Brat-NHL fits into the picture remained unclear. Several conclusions can be drawn from our results. First, our binding assays reconcile the published results on RNA-binding by Pum-HD and Brat-NHL. Pum-HD was shown to bind both Box A and Box B, binding with high affinity to Box B and lower affinity to Box A (16). Box A, however, is the binding site of Brat-NHL and in fact binding by Pum-HD was proposed to enhance the Brat-NHL-RNA interaction, putatively by unfolding the secondary structure of the RNA (22). Our binding assays show that the secondary binding of Pum-HD to Box A is of lower affinity than the binding of Brat-NHL (∼2 μM K_d_ comparing to K_d_ of ∼1.2 μM), so Box A is preferentially bound by Brat-NHL. Furthermore, we show that incubating the RNA with Pum-HD does not enhance the Brat-NHL-RNA interaction. The previously reported enhancement was based purely on EMSA experiments, which are not quantitative equilibrium assays. Moreover, the reported effect was very mild and likely in the error range of the method. We further confirm that Pum-HD indeed unfolds the secondary structure of the RNA. However, we find no evidence that it enhances the Brat-NHL-RNA interaction and find no reasons to assume so, particularly considering that the reported Brat-NHL binding site lies predominantly outside of the secondary structure of the RNA (Figure 1A). In conclusion, we show that Pum-HD has no significant effect on the binding of Brat-NHL to the RNA, suggesting that Pum-HD and Brat-NHL recognize the *hunchback* mRNA neither competitively nor cooperatively, but independently.

Another option for cooperativity in *hunchback* mRNA recognition between the Pumilio-Nanos moiety and Brat-NHL could be Nanos-mediated cooperativity. Thus, we thoroughly investigated the relationship between Nanos-ZnF and Brat-NHL. First, we do not observe any signs of interaction between the domains *in vitro* in the absence of RNA. However, as illustrated by the Nanos-ZnF-Pum-HD interaction seen only in the ternary RNA complex (15), additional protein-protein interactions might be induced by RNA-binding. We then tested for a potential Nanos-ZnF-Brat-NHL interaction induced by *hb* complex formation by comparing NMR spectra of Brat-NHL in its free form, RNA-bound form and Brat-NHL in the *hb* complex. If an additional interaction induced by RNA-binding exists, we would expect to observe chemical shift perturbations of Brat-NHL resonances upon *hb* complex formation. However, the NMR spectra of Brat-NHL bound to NRE2 RNA and of Brat-NHL in the *hb* complex look identical.

The last conceivable possibility for cooperativity between the Pumilio-Nanos pair and Brat-NHL could be via the RNA. Similar mechanism as proposed earlier for Pum-HD and Brat-NHL could exist – Pumilio-Nanos moiety could alter the structure of NRE2 RNA, thus making the RNA more accessible for Brat-NHL. Such a cooperativity may not lead to any protein-protein interaction. However, it should manifest as a noticeable increase in the affinity of the Brat-NHL to the Pum-HD-Nanos-ZnF-bound NRE2 RNA. For this reason we performed ITC measurements with Brat-NHL and preformed Pum-HD-Nanos-ZnF-RNA complex. The results show that Brat-NHL binds the Pum-HD-Nanos-ZnF-bound NRE2 RNA with the same affinity as free NRE2 RNA. Collectively, our results dismiss the possibility of cooperativity in RNA-binding between Brat-NHL and Pum-HD with Nanos-ZnF.

The binding sites of Brat-NHL and Pum-HD with Nanos-ZnF are separated by four nucleotides, so if there is no cooperativity in the RNA-binding and Pum-HD unfolds the secondary structure of the RNA, there should be no constraints on these four nucleotides. The unconstrained nucleotides should then give the *hb* complex flexibility allowing free movement of the Pum-HD-Nanos-ZnF and Brat-NHL moieties. We therefore derived an atomistic conformationally free ensemble from unbiased all-atom MD simulations, which quantitatively agreed with the SAXS data, suggesting that the *hb* complex is indeed largely flexible. Our NMR relaxation measurements then confirm the flexibility, as we see no difference between the rotational correlation time of Brat-NHL in its free form and in the *hb* complex.

It is well established that disordered proteins or protein complexes play key roles in cellular function and disease, however, characterizing such systems at atomic detail remains challenging. We anticipate that the integration of various structural and biophysical data with atomistic simulations, as carried out here, will be mandatory to develop mechanistic understanding of disorder in biology.

In summary our data clearly show that the RBD of Brat, and RBDs of Pumilio and Nanos, interact with *hunchback* RNA independently. Considering that the proteins are able to suppress translation independently (22,23,81), this could indicate that the suppression of *hunchback* mRNA by Brat is functionally separate of the suppression by Pumilio with Nanos. The translation of the maternal *hunchback* mRNA is suppressed from early embryogenesis to maternal-to-zygotic transition (MZT) (82). During MZT the maternally supplied mRNA is degraded by maternal and zygotic activities, so the embryo switches from maternal to zygotic genome (83). mRNA decay is often coupled to translation suppression and in fact Brat-mediated translation suppression was shown to control the maternal activity in RNA decay during MZT (84). Brat is distributed uniformly throughout the embryo (14), but Nanos is expressed in a gradient (4). If *hunchback* mRNA translation suppression by Brat and by the Pumilio-Nanos pair are separate, the suppression by Pumilio and Nanos might serve to establish the gradient of Hunchback in early embryogenesis, while the suppression by Brat might serve to control the maternal activity in RNA decay during MZT.

Considering all of these findings, there are two major questions to answer in order to gain a definitive and full understanding of the mechanism of *hunchback* mRNA translation suppression. First, it is crucial to extend this biophysical investigation ultimately to full-length proteins and RNA covering both NREs to test if Brat and Pumilio with Nanos still bind the RNA independently, especially considering that Brat has a dimeric coiled-coil domain (85). Secondly, functional experiments are necessary to decipher if the suppression activities of Brat and Pumilio with Nanos are separate. The suppression of *hunchback* mRNA translation is one of the most-well studied cases of localized translation suppression during development, so elucidating its mechanism fully will not only provide insights into the function of the specific *trans*-acting molecules, but will also give clues about general mechanisms of localized translation suppression during development.

## ACKNOWLEDGEMENT

We thank the EMBL/DESY Hamburg PETRA-III (P12 beamline), and the ILL (D22 beamline) local contact for support (Anne Martel). This work was supported by an Emmy-Noether Fellowship (HE 7291_1) and the Priority Program SPP1935 of the Deutsche Forschungsgemeinschaft (DFG) and the EMBL. We also thank Inga Loedige for providing some of the constructs used and for fruitful discussion.

