## Supplementary Material for "Structure and dynamics of the quaternary *hunchback* mRNA translation repression complex"

### SUPPLEMENTARY DATA

Supplementary Data are available at NAR online.

| Experiment | Cell | Syringe | Injections | Injection size |
|---|---|---|---|---|
| Figure 1A | 20 μM NRE2 RNA | 400 μM Pum-HD | 16 | 2.5 (0.4) μl |
| Figure 1B | 25 μM NRE2 RNA | 375 μM Brat-NHL | 13 | 3 (0.4) μl |
| Figure 1C | 24 μM NRE2 RNA + Pum-HD | 375 μM Brat-NHL | 13 | 3 (0.4) μl |
| Figure X | 10 μM NRE2 RNA + Pum-HD + Nanos-ZnF | 200 μM Brat-NHL | 13 | 3 (0.4) μl |

**Table S1.** Details of the ITC titrations. The bracketed number in the injection size column states the size of the initial injection.

| | | Concentrations tested (mg/ml) | | |
|---|---|---|---|---|
| Screen | Supplier | 4°C | 20° C | Details |
| Classics | Qiagen | 1.5, 3.0, 5.13, 10.0 | 1.5, 3.0, 5.13, 10.0 | - |
| JCSG+ | Molecular Dimensions | 1.5, 3.0, 5.13, 10.0 | 1.5, 3.0, 5.13, 10.0 | - |
| PACT | Molecular Dimensions | 1.5, 3.0, 5.13 | 1.5, 3.0, 5.13 | - |
| PEGS | Qiagen | 1.5, 3.0, 5.13 | 1.5, 3.0, 5.13 | - |
| Wizzard I+II | Rigaku | 1.5, 3.0, 5.13, 10.0 | 1.5, 3.0, 5.13, 10.0 | - |
| Natrix | Hampton Research | 1.5, 3.0, 5.13, 10.0 | 1.5, 3.0, 5.13, 10.0 | - |
| Nucliex | Qiagen | 1.5, 3.0, 5.13, 10.0 | 1.5, 3.0, 5.13, 10.0 | - |
| MPD | Qiagen | 1.63, 2.13, 2.63, 3.13, 3.63, 4.13, 4.63, 5.13 | - | - |
| $AmSO_4$ | Qiagen | 1.5, 3.0, 5.0 | 3.0, 5.0 | - |
| MIDAS | Molecular Dimensions | 1.5, 3.0, 5.0 | 1.5, 3.0, 5.0 | - |
| Morpheus | Molecular Dimensionns | 1.5, 3.0, 5.0 | 1.5, 3.0, 5.0 | - |
| PEGs II | Qiagen | 1.5, 3.0, 5.0 | 3.0, 5.0 | - |
| SaltRX | Hampton Research | 1.5, 3.0, 5.0 | 3.0, 5.0 | - |
| The BCS screen | Molecular Dimensions | 3.0 | 3.0 | - |
| $AmSO_4$ vs. PEG400 | Custom | 1.5, 2, 3, 4 | - | 0-2.2 M $AmSO_4$, 0-14% PEG, 0.1 M MES pH 6.5 |
| $AmSO_4$ vs. PEG3350 | Custom | 1.5, 2, 3, 4 | - | 0-2.2 M $AmSO_4$, 5-19% PEG, 0.1 M MES pH 5.6 |
| $AmSO_4$ vs. pH | Custom | 1.5, 2, 3, 4 | - | 0-2.2 M $AmSO_4$, 5% PEG400, 0.1 M MES pH 5.5-6.7 |
| $SrCl_2$ vs. $Li_2SO_4$ | Custom | 3 | 3 | 0.005-0.2 M $SrCl_2$, 0.1-1.9 $Li_2SO_4$, 0.1 M Na-Acetate pH 4.6 |
| pH vs. $Li_2SO_4$ | Custom | 3 | 3 | 0.1 M $SrCl_2$, 0.1-1.9 $Li_2SO_4$, 0.1 M Na-Acetate pH 3.8-5.2 |
| pH vs. PEG mix | Custom | 2.5, 5 | 2.5, 5 | 0.1 M Tris, BICINE pH 8.2-8.9, 10-32% v/v Ethylene glycol, 5-16% w/v PEG 800 |
| pH vs. PEG mix | Custom | 2.5, 5 | 2.5, 5 | 0.1 M Na-HEPES; MOPS pH 7.2-7.9, 10-32% v/v Ethylene glycol, 5-16% w/v PEG 800 |
| pH vs. PEG mix | Custom | 2.5, 5 | 2.5, 5 | 0.1 M Tris, BICINE pH 8.2-8.9, 10-32% v/v Ethylene glycol, 5-16% w/v PEG 800, 0.03 M $MgCl_2$, 0.03 M $CaCl_2$ |

**Table S2.** Crystallization trials of *hb* complex. The trials were set up by mixing 100 nl protein solution and 100 nl of the mother liquor. Hanging drop set up and vapour diffusion methods were used.

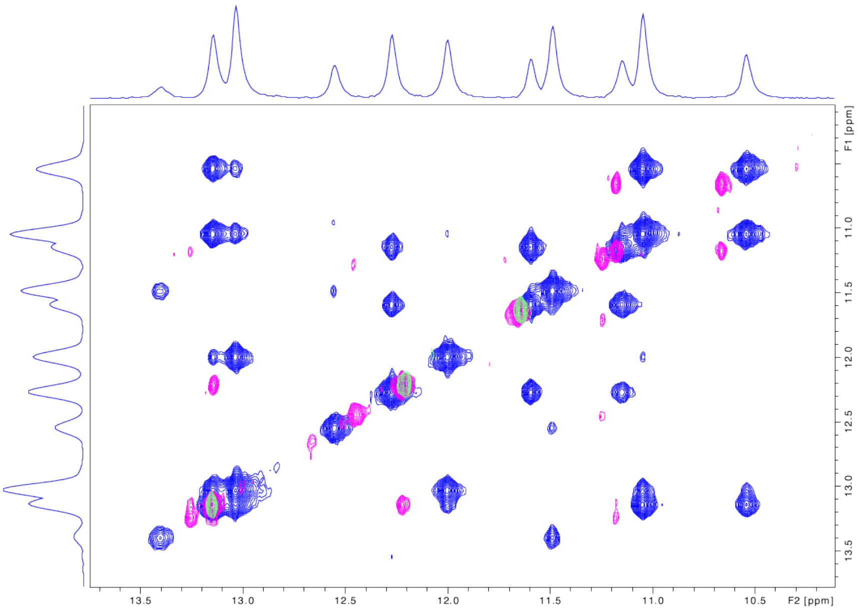


**Figure S 1.** Purified Pum-HD unfolds *hunchback* mRNA. The figure shows an overlay of the imino region of the $^{1}H$, $^{1}H$ NOESY spectra of *hunchback* mRNA. The RNA consists of directly connected NRE1 and NRE2. In blue is the spectrum of the RNA at 5 °C, in purple is the spectrum of the RNA at 20 °C and in green is the spectrum of the RNA with an equimolar amount of Pum-HD at 20 °C. The purple and blue cross-peaks for example at 13.2,12.2 ppm (F1,F2) or at 11.2,10.6 pmm (F1,F2) illustrate how hydrogen bonds formed in base pairing disappear upon the addition of Pum-HD.

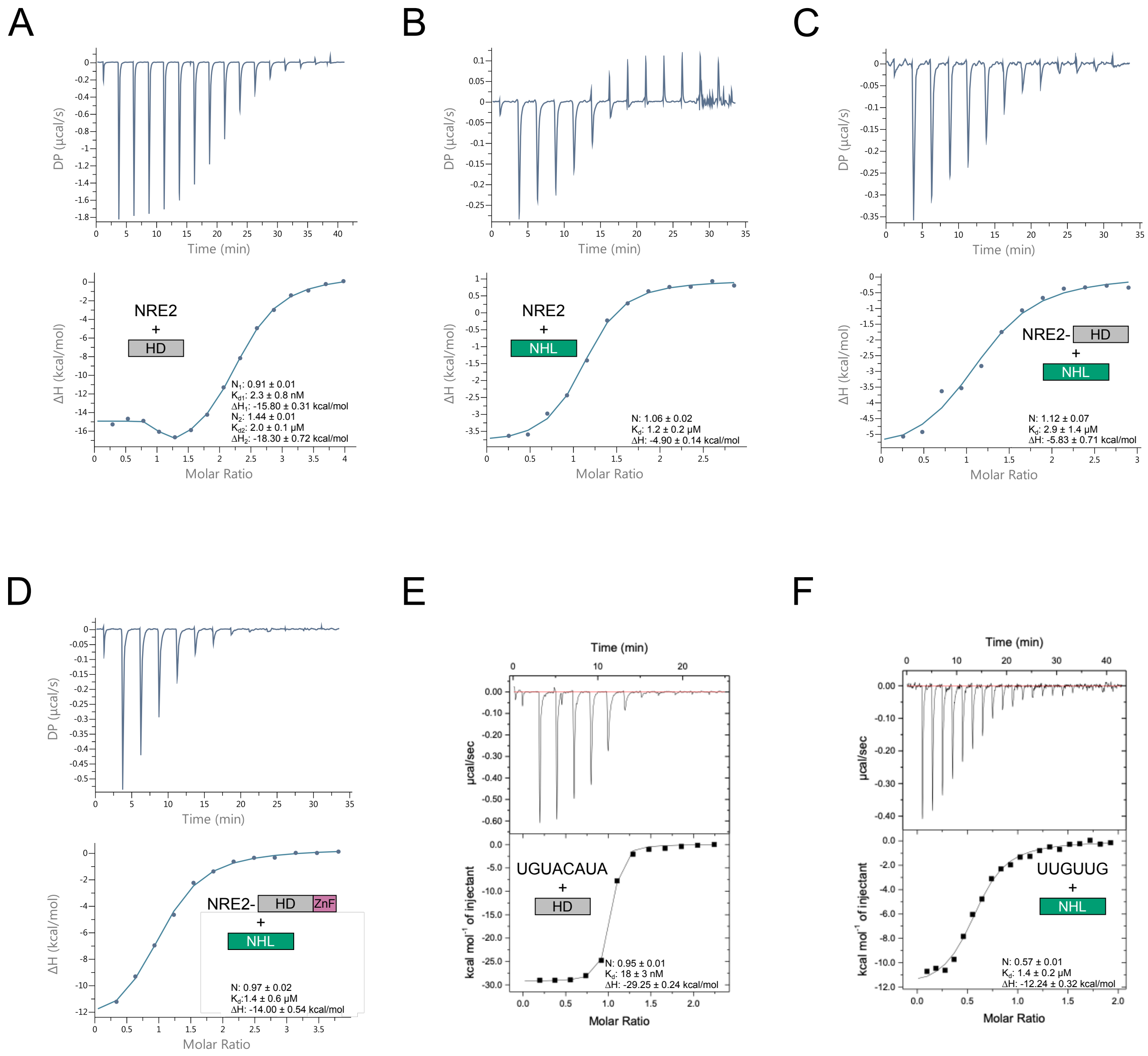


**Figure S2.** Full results of the ITC measurements. The panels correspond to measurements of the following interactions: **(A)** Pum-HD-NRE2 RNA interaction, **(B)** Brat-NHL-NRE2 RNA interaction, **(C)** the interaction of Pum-HD-bound NRE2 RNA with Brat-NHL, **(D)** the interaction of Brat-NHL with the Pum-HD-Nanos-ZnF-NRE2 RNA complex, **(E)** the interaction of Pum-HD with an RNA encompassing its binding site (full sequence of the RNA given in main figure 1) and **(F)** the interaction of Brat-NHL with an RNA encompassing its binding site (full sequence of the RNA given in main figure 1).

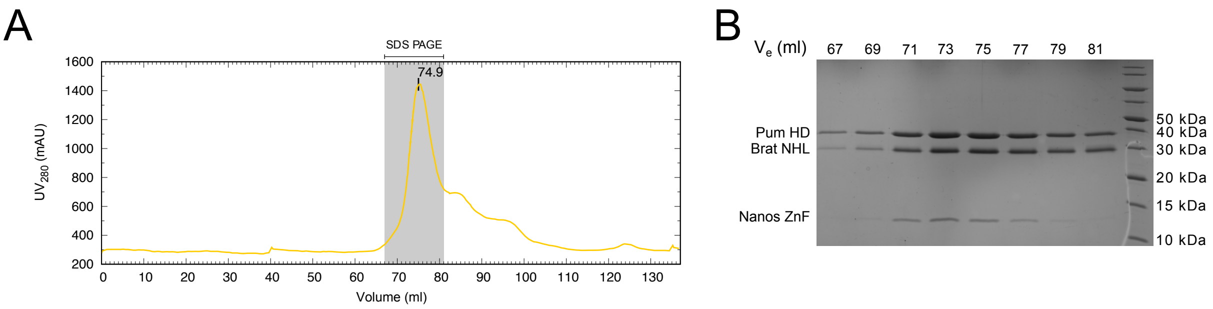


C

| | | |
|---|---|---|
| Calculated MW | Brat NHL | 31 925.45 Da |
| | Nanos ZnF | 10 550.10 Da |
| | Pum HD | 38 372.09 Da |
| | NRE2 RNA | 6 950.40 Da |
| | Total | 87 798.04 Da |
| SEC | $V_e$ | 74.9 ml |
| | MW | ~ 77 000 Da |
| MALLS | Mw | 84 935 ± 1676 Da |
| | Mw/Mn (polydispersity) | 1 ± 0.03 |

**Figure S3.** Reconstitution of the *hb* complex. To reconstitute the *hb* complex purified Pum-HD, Nanos-ZnF and Brat-NHL were mixed with NRE2 RNA as described in Material and Methods. SEC was then used to purify the complex and its identity was confirmed by SDS-PAGE, UV absorption and MALLS. **(A)** SEC chromatogram of *hb* complex purification. SEC was done using HiLoad 16/600 Superdex 200 pg column (GE Healthcare). The grey shaded area indicates the analysed fractions of the *hb* complex peak. **(B)** SDS-PAGE of the selected fractions from SEC. The gel used was 15% AA. **(C)** Comparison of expected and experimentally determined molecular weights of the *hb* complex. MALLS was done in an online SEC-MALLS setup at P13 beamline at DESY, Hamburg, Germany.

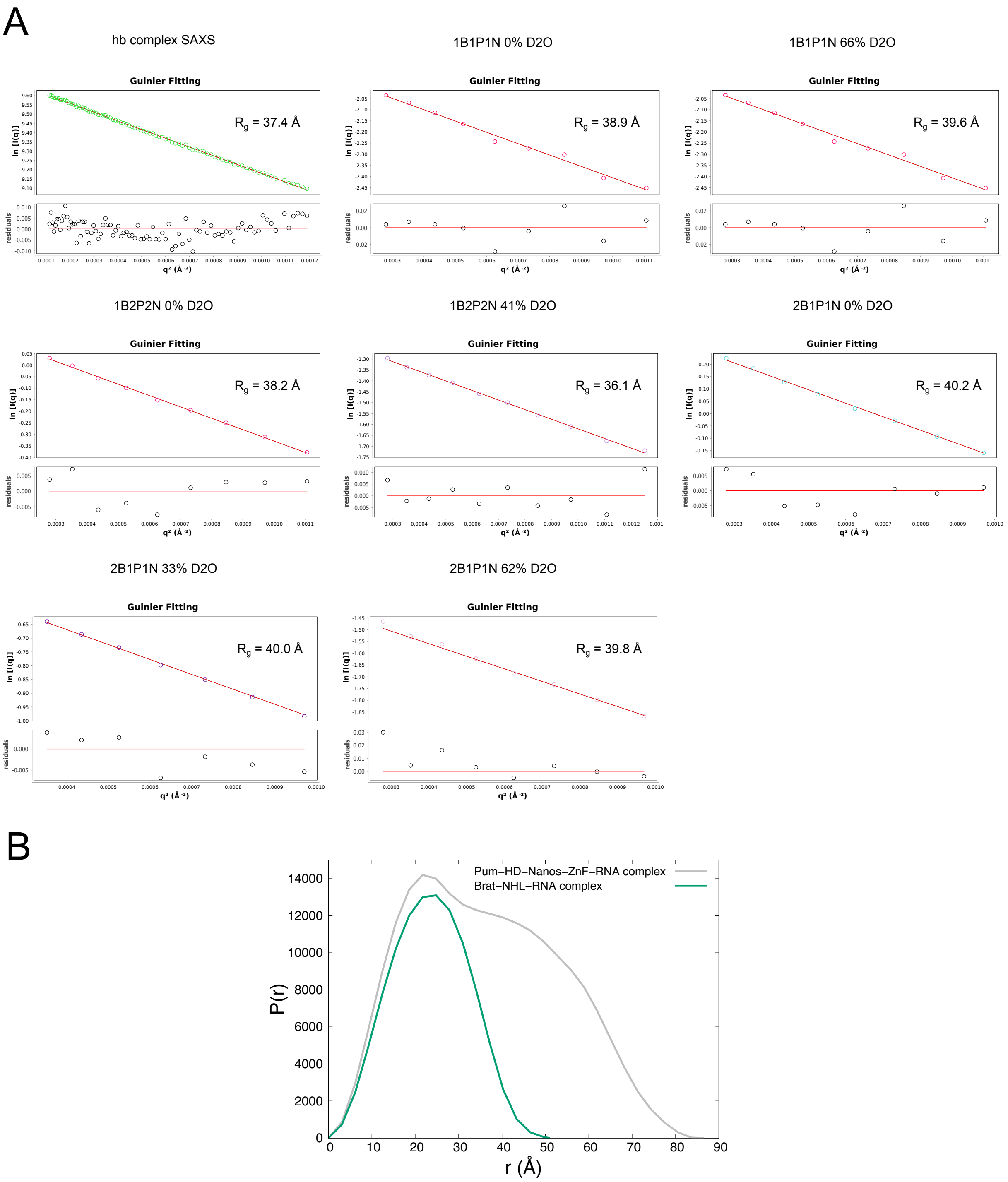


**Figure S4**. Characterization of *hb* complex by small angle scattering (SAS). **(A)** Guinier fit from the individual scattering curves. All fits obey $q*R_g < 1.3$. **(B)** Distance distribution function (P(r)) of Brat-NHL-RNA complex and Pum-HD-Nanos-ZnF-RNA complex. P(r) were calculated from the available high-resolution structures of Brat-NHL-RNA complex (PDB ID: 4zlr) (1) and Pum HD-Nanos ZnF-RNA complex (PDB ID: 5kl1) (2) using ScÅtter (3).

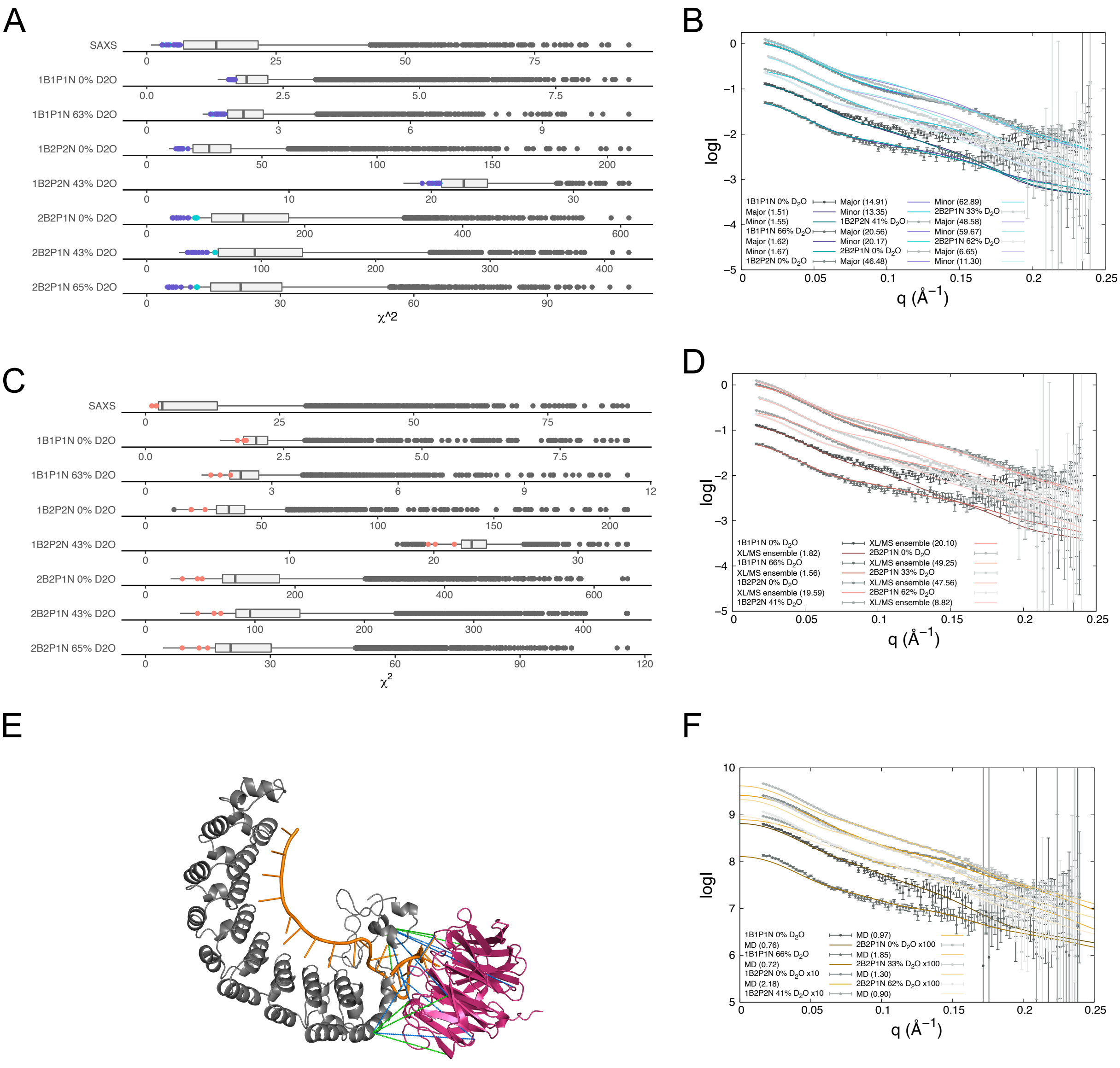


**Figure S5.** Fits of *hb* complex models to the experimental data. **(A)** Distribution of $\chi^2$ values of fits of the pool of models generated to model *hb* complex based on SAS data. To model the *hb* complex 5055 random models were generated and theoretical scattering curves of each model at each condition were then back-calculated and fitted against the experimental data using CRYSOL and CRYSON. The best fitting models were selected as the models in the top 0.22 quantile of $\chi^2$ values of fits of each curve and are highlighted in the distribution as either the purple or cyan dots according to whether they fall into the major or minor cluster. **(B)** Fits of the major and minor cluster of models to the experimental small-angle neutron scattering (SANS) data. For each cluster a fit of a representative model is shown. A representative model of the cluster was selected as the model closest to the mean coordinates of the aligned cluster. **(C)** Distribution of $\chi^2$ values of fits of the pool of models generated to model *hb* complex based on SAS and XL/MS data. To model the hb complex 4572 models were generated and theoretical scattering curves of each model at each condition were then back-calculated and fitted against the experimental data using CRYSOL and CRYSON. The models in the top 0.30 quantile of $\chi^2$ values of each curve were selected and then this pool was further restricted to only the models in the top 0.10 quantile of the lowest distance restraint violation energy. The selected ensemble is highlighted in the distribution as magenta dots. **(D)** Fit of the selected ensemble of *hb* complex models generated by modelling using SAS and XL/MS data to the experimental SANS data. The fit of the ensemble to the experimental data is illustrated by the fit of the model closest to the mean coordinates of the aligned ensemble. **(E)** Satisfaction of the XL/MS data for a representative model of *hb* complex. The representative model is the model with the lowest distance restraint violation energy from the selected ensemble of models obtained by modelling using SAS and XL/MS data. Green lines depict satisfied cross-links, whereas blue lines show violated cross-links. **(F)** Fit of the ensemble of *hb* complex models obtained by all-atom MD simulations. The curve fitted in each condition is a back-calculated scattering curve of frames from the time interval between 30 and 110 ns from the all-atom MD simulations. For clarity, curves for 1B2P2N and 2B2P1N were offset along the Y-axis by multiplying the curves with factors of 10 and 100, respectively. To avoid any influence by aggregation in the SANS data, the fits were restricted to q>0.07Å (2B2P1N) or to q>0.05Å (all other sets). The number in the brackets always reports the values of $\chi^2$ values of the fit.

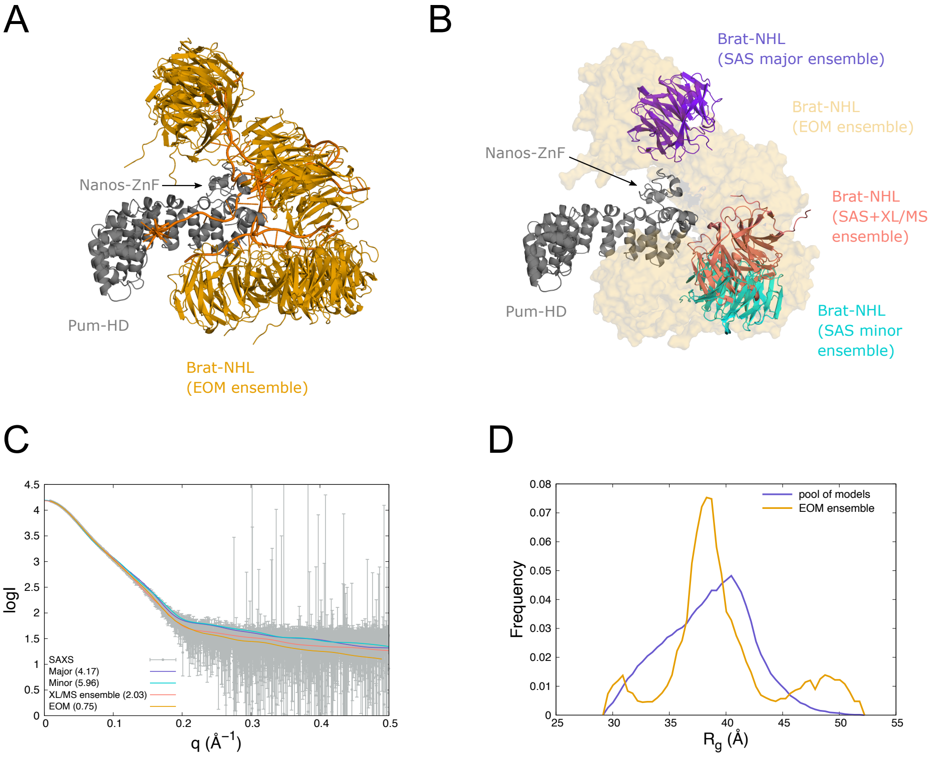


**Figure S6.** Modelling the *hb* complex by Ensemble Optimization Method (EOM) (4). **(A)** Ensemble of *hb* complex models obtained from EOM using the SAXS curve of the complex. EOM used the starting pool of models generated during rigid-body modelling to search for an ensemble of models to fit the SAXS data as a mixture. Seven models of the best fitting ensemble are shown. **(B)** Overlay of *hb* complex models. Representative models of ensembles from rigid-body are superimposed on the EOM ensemble from (A) with Brat-NHL in semi-transparent surface representation. All models in the ensembles are always superimposed on Pum-HD and Nanos-ZnF. **(C)** Fits of back-calculated scattering curves of representative models of *hb* complex ensembles to the experimental SAXS curve. 'Minor', 'Major', and 'XL/MS ensemble' denotes the representative models of the ensembles obtained from the rigid-body modelling. Numbers in the brackets state the $\chi^2$ value of the fit. **(D)** $R_g$ distribution plot from EOM of the *hb* complex. The plot shows the distribution of the $R_g$ values of the initial pool of models (generated during rigid-body modelling), and the $R_g$ distribution of the ensemble of models selected by EOM to fit the SAXS data (EOM ensemble). The final ensemble has 7 models with an $R_{flex}$ of 88.3% ($R_{flex}$ of the pool 89.5%) and $R_\sigma$ of 5.26.


1. Loedige, I., Jakob, L., Treiber, T., Ray, D., Stotz, M., Treiber, N., Hennig, J., Cook, K.B., Morris, Q., Hughes, T.R. *et al.* (2015) The Crystal Structure of the NHL Domain in Complex with RNA Reveals the Molecular Basis of Drosophila Brain-Tumor-Mediated Gene Regulation. *Cell Rep*, **13**, 1206-1220.
2. Weidmann, C.A., Qiu, C., Arvola, R.M., Lou, T.F., Killingsworth, J., Campbell, Z.T., Tanaka Hall, T.M. and Goldstrohm, A.C. (2016) Drosophila Nanos acts as a molecular clamp that modulates the RNA-binding and repression activities of Pumilio. *Elife*, **5**.
3. Forster, S., Apostol, L. and Bras, W. (2010) Scatter: software for the analysis of nano- and mesoscale small-angle scattering. *Journal of Applied Crystallography*, **43**, 639-646.
4. Tria, G., Mertens, H.D., Kachala, M. and Svergun, D.I. (2015) Advanced ensemble modelling of flexible macromolecules using X-ray solution scattering. *IUCrJ*, **2**, 207-217.
